# Mapping Core Components of Membrane-less Organelles in Living Cells by Phase-APEX2 Proximity Labeling

**DOI:** 10.64898/2025.12.17.694854

**Authors:** Zhuo Chen, Yongzuo Chen, Huarun Ding, Miaodan Huang, Weijie Chen, Bin Fu, Mingchao Wang, Lingqiang Zhang, Wei Qin, Pilong Li

**Affiliations:** Tsinghua-Peking Center for Life Sciences, Beijing Frontier Research Center for Biological Structure, State Key Laboratory of Membrane Biology, Tsinghua University, Beijing, China; School of Life Sciences, Tsinghua University, Beijing, China; School of Pharmaceutical Sciences, MOE Key Laboratory of Bioorganic Phosphorus Chemistry & Chemical Biology, Tsinghua University, Beijing, China; State Key Laboratory of Medical Proteomics, National Center for Protein Sciences (Beijing), Beijing Institute of Lifeomics, Beijing 102206, China

## Abstract

Membrane-less organelles (MLOs), formed via liquid-liquid phase separation, are crucial for organization of biomacromolecules and cellular processes, yet their dynamic nature challenges compositional analysis. Conventional proximity labeling methods lack the specificity to overcome high background and systematic biases inherent in studying these condensates. Here we introduce Phase-APEX2, a three-input AND-gate strategy that dramatically enhances labeling specificity. By integrating a split-APEX2 system with a light-controlled H_2_O_2_ generator, we developed Phase-APEX2 platforms for two application contexts: Phase-APEX2-MS for proteomics and Phase-APEX2-seq for genomics. Our analyses revealed not only the core interactomes of the nucleolus and stress granules with high accuracy but also uncovered distinct “molecular grammars” governing their assembly and a surprising link between nucleolar dynamics and the transcriptional regulation of mitosis-related genes. This highly specific and adaptable platform opens new avenues for exploring the composition and function of diverse biomolecular condensates.

## Introduction

MLOs, such as the nucleolus^1–3^ and stress granules^4–8^, are dynamic mesoscale structures formed through the liquid-liquid phase separation (LLPS) principle. As such, they continuously exchange their compo-nents with the surroundings. Traditional methods (such as differential centrifugation and particle sorting) cannot isolate them cleanly without contamination from other cellular components^9,10^. Proximity labeling (PL) techniques, which are able to map subcellular components in living cells, have recently emerged as powerful tools compared to traditional biochemical approaches^11,12^.

However, the unique properties of MLOs pose several challenges for conventional proximity labeling technologies (Figure 1A). First, the composition of these dynamic condensates changes rapidly in response to cellular stimuli, demanding techniques with high temporal resolution (minutes or even seconds)^9,13–15^. Second, many MLO components exhibit multi-localization patterns^16^. For example, numerous nucleolar proteins also reside at the cell membrane^16–19^, or shuttle between the nucleolus and stress granules during stress conditions^20^. Multi-localization patterns of MLO proteins make it difficult to identify a truly specific marker to serve as a bait for fusion with a labeling enzyme. Third, MLO components face a concentration-volume trade-off^14,21^. Although MLOs concentrate macromolecules, the enrichment of MLO proteins in cellular contexts is often modest (e.g., ∼10-fold for G3BP1 in stress granules^6,7^). Moreover, MLOs are typically small (100-1000 nm in diameter^22^), occupying only a minor fraction (<1%) of the cellular volume. Compounding this issue, studies indicate that for many canonical components, the majority of the protein pool (>80%) remains in the dilute phase, with only a small fraction (<20%) partitioned into the condensates^14,21^, which leads to high background catalytic signals outside the MLOs.

**Figure 1.**
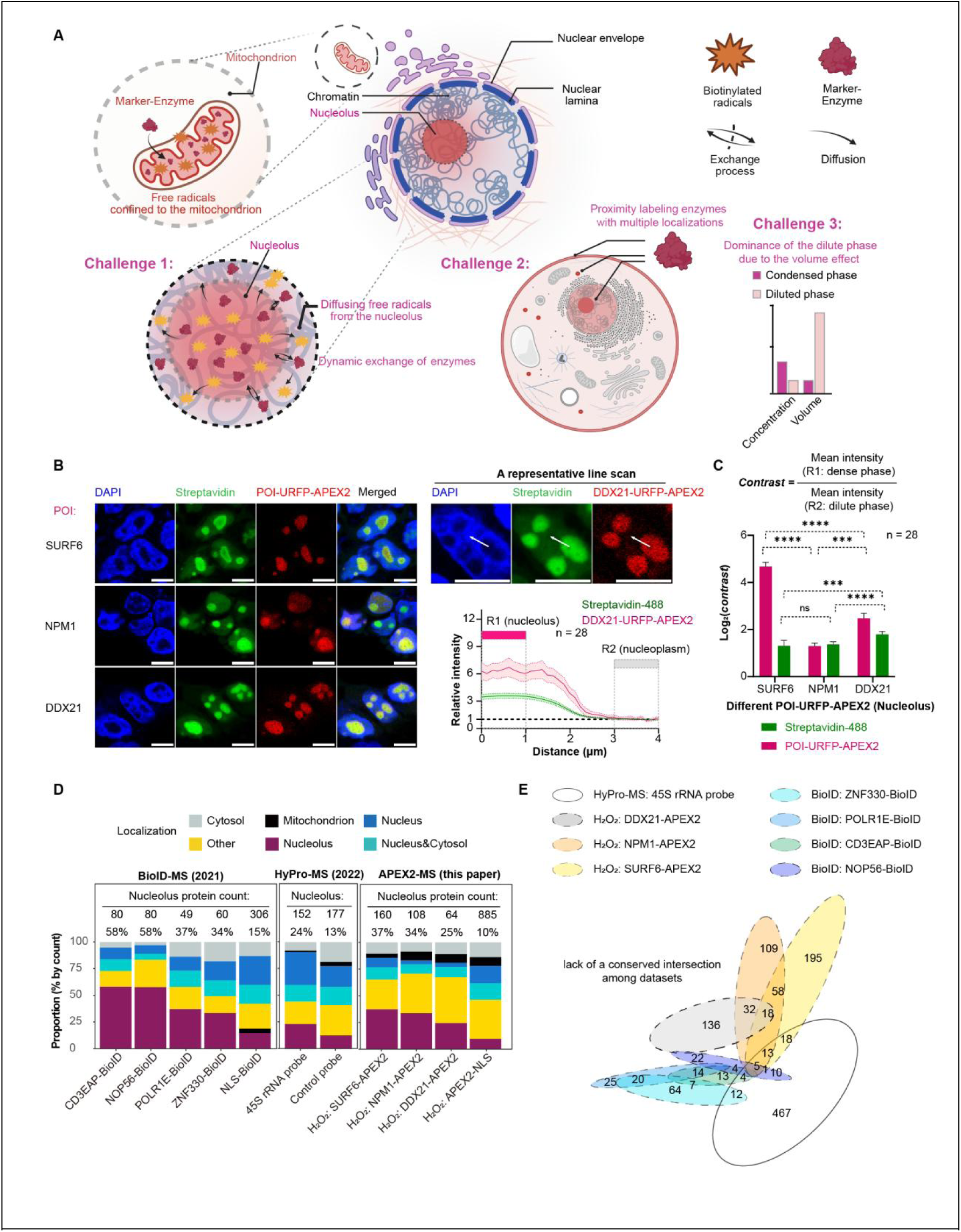
Conventional proximity labeling strategies show limited utility in membraneless organelles. (A) Schematic illustrating the 3 major challenges when using common proximity labeling (PL) techniques to analyze MLOs. (B-C) Confocal microscopy of 293T cells used for PL experiments (B) and fluorescence contrast comparison (C) across POI groups. Data are mean ± 95% CI (n = 28); Kruskal-Wallis test; ns, not significant, ***adj. p (BH corrected) < 0.001, ****adj. p < 0.0001. (D) Specificity analysis of published datasets (BioID-MS, HyPro-MS) and data from our APEX2-MS method in the nucleolus system. Euler diagram of final protein lists obtained from 8 different PL datasets. Scale bars, 10 μm.

The existing proximity labeling toolkit is inadequate to overcome these challenges. Conventional single-bait strategies, including APEX2^23^, TurboID^24^, and other photo-activated proximity labeling techniques employing single photosensitizers (μMap^25,26^, MultiMap^27^) or enzymes^28–30^, rely critically on the precise subcellular localization of the bait-fusion protein. This strategy is fundamentally limited by the multi-localization and concentration-volume trade-offs (challenges 2 and 3 in Figure 1A), making specific MLO targeting problematic. Even when an appropriate bait protein is identified, these single-input approaches are ill-suited for complex applications such as investigating MLO subpopulations or inter-organelle contact sites. To improve targeting flexibility, dual-bait strategies including split-APEX2^31^ and split-TurboID^32^ were developed. However, these methods remain constrained by their requirement for exogenous substrate addition (*e.g.*, hydrogen peroxide or biotin), which limits the temporal resolution (challenge 1 in Figure 1A). Recent studies have described several light-activated or chemically-inducible dual-bait proximity labeling techniques^33,34^, but their specificity and general applicability for profiling diverse MLO components (nucleic acids/proteins) have not been comprehensively evaluated.

Here, we developed Phase-APEX2, a three-input AND-gate photocatalytic strategy that overcomes these limitations. Phase-APEX2 integrates a split-APEX2 system with a genetically encoded, light-controlled H_2_O_2_ generator to create a 3-input AND-gate logic system. This design leverages the co-enrichment of three components to amplify the labeling rate difference between the condensed and dilute phases, thus minimizing off-target effects while preserving APEX2’s rapid kinetics. In nucleoli and stress granules, Phase-APEX2-MS, coupled with a novel 2D data filtering model, identified core MLO proteins within high-density protein-protein interaction regions that were not detected by other 1-input/2-input methods, increasing the area under the receiver operating characteristic curve (AUROC) to 0.8-0.9. Notably, we discovered that the core proteins of stress granules and nucleoli exhibit fundamentally distinct molecular grammar. Following protocol optimization, we integrated Phase-APEX2 with next-generation sequencing (NGS) to develop Phase-APEX2-seq. This approach successfully identified nucleolus-associated genomic regions with high accuracy and revealed that specific gene clusters are strongly associated with cell division, suggesting that these regions may regulate cell cycle progression in synchrony with nucleolar dynamics.

## Results

### Conventional single-marker proximity labeling lacks specificity for MLOs

To assess whether single-bait proximity labeling provides sufficient specificity for MLO profiling, we systematically evaluated its performance in both nucleoli and stress granules of 293T cells. For nucleolar targeting, we fused three distinct marker proteins (SURF6, NPM1, and DDX21)^35^ to APEX2 respectively. Although these fusion proteins exhibited enrichment within nucleoli, they all generated visible biotinylation signals emanating from the surrounding nucleoplasm (Figure 1B). While bait protein enrichment varied considerably (2- to 24-fold), the resulting biotinylated product enrichment remained consistently low (∼2-to 4-fold) (Figure 1C). Reducing the labeling time from 120 s to 30 s failed to enhance specificity, with enrichment values persistently below 4-fold (Figure S1G).

A comparable deficiency in specificity was observed in the stress granule (SG) system using G3BP1 and TIA1 as markers. Fusion protein enrichment within SGs (6- to 8-fold) significantly exceeded the corresponding catalytic product enrichment (∼4-fold) (Figure S1A-C). More strikingly, TIA1-APEX2 exhibited anomalous catalytic activity despite the protein’s predominantly cytoplasmic localization, its catalytic signal was aberrantly concentrated within the nucleus (Figure S1D). Quantitative analysis of nucleus-to-cytoplasm (Nuc/Cyto) fluorescence ratios confirmed that TIA1-APEX2’s catalytic signal was always enriched in the nucleus, directly opposing the bait’s subcellular distribution (Figure S1E-F) and implying strong off-target catalysis in the nucleus.

To corroborate these imaging findings with proteomic evidence, we compared nucleolar proximity labeling results across published BioID^36^ and HyPro-MS^37^ datasets alongside our own proteomic data generated using conventional APEX2-MS protocols^10^ (Figure 1D). The proportion of known nucleolar proteins identified varied substantially, ranging from 34–58% for BioID to 25–37% for APEX2, and merely 24% for HyPro-MS. In two-dimensional scatter plots, most of the known nucleolar proteins failed to form discrete clusters separate from background proteins and were distributed across different regions, indicating that no single threshold criterion can reliably generate a specific proteome across different datasets (Figure S1H). Euler diagram analysis further substantiated this methodological discordance, revealing the absence of a conserved core set of overlapping proteins and highlighting significant inter-technique variability (Figure 1E).

Collectively, these complementary imaging and proteomic analyses demonstrate that conventional single-marker proximity labeling strategies suffer from inadequate specificity and substantial systematic errors when applied to MLOs.

### A split-APEX2 strategy improves labeling specificity

To enhance the specificity of proximity labeling, we developed a two-input AND-gate using a split-APEX2 system^31^ (Figure 2A, Module 1). We hypothesized that the high local concentration within MLOs would facilitate the reconstitution of split-APEX2 fragments, thereby promoting preferential catalytic activity specifically within the condensate. Although split-APEX2 typically requires heme supplementation^31^ *in vitro*, we discovered that endogenous heme^38–40^ in cellular contexts was adequate to sustain proximity labeling activity (Figure S2C). The system functioned as a stringent AND-gate, as evidenced by the complete absence of catalytic activity when either fragment was expressed individually in the stress granule system (Figure S2B). Notably, co-expression of G3BP1-AP and TIA1-EX in SGs yielded a catalytic signal with ∼2-fold greater specificity than single-bait methods and eliminated the aberrant nuclear catalysis of TIA1-APEX2 (Figure 2B, S2A). A systematic evaluation of nine split-APEX2 pair combinations (generated by fusing three nucleolar markers to either the N- or C-terminal fragments of APEX2, resulting in 3×3 combinations) in the nucleolus confirmed an average 1.5- to 2-fold specificity increase over single-marker approaches (Figure S2D-E). These results validate the split-APEX2 strategy as an effective 2-input AND-gate that improves labeling selectivity for MLOs.

**Figure 2.**
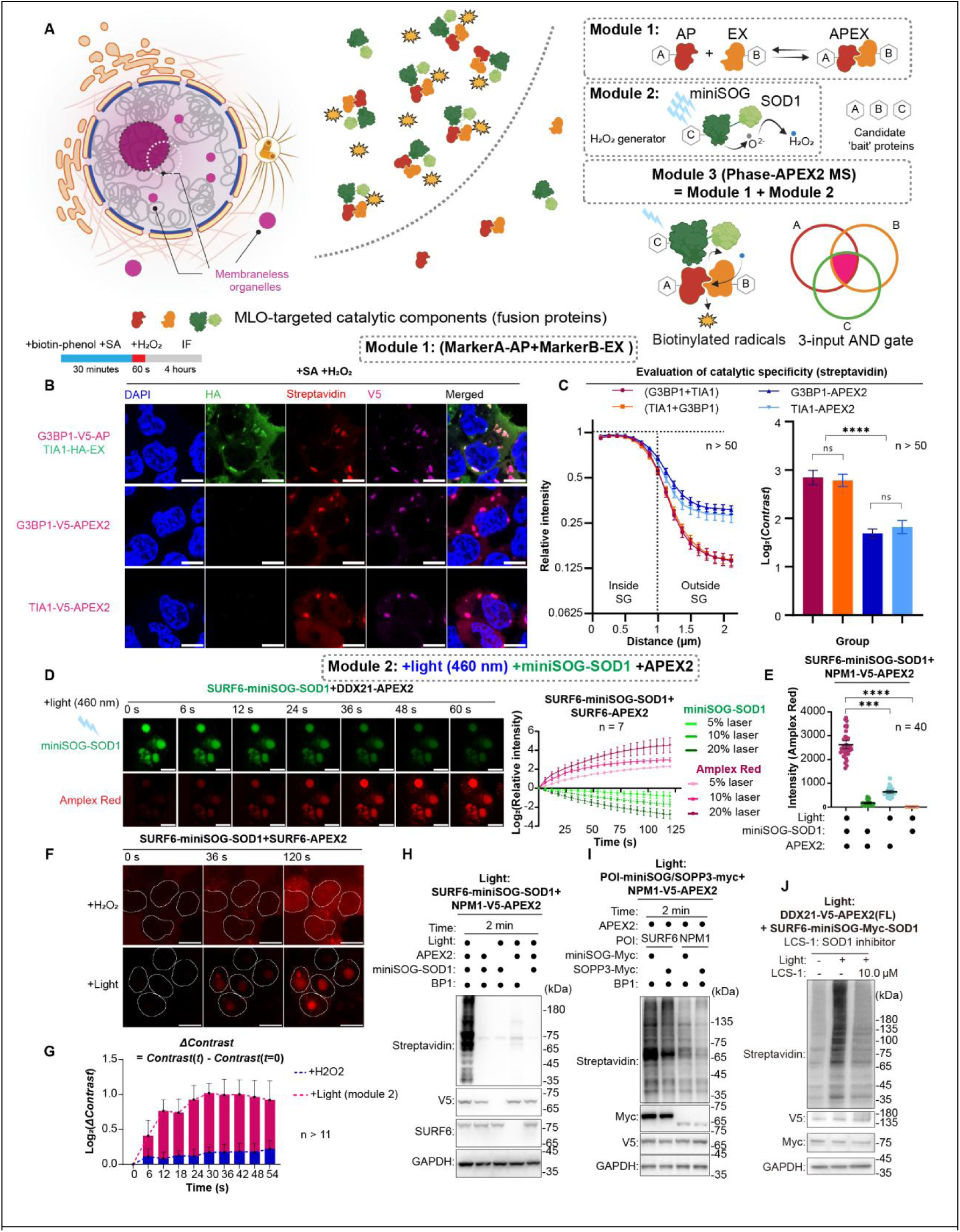
Enhancing the spatial selectivity of proximity labeling with split-APEX2 and an *in situ* H_2_O_2_ generator. (A) Schematics of module 1 (split-APEX2) and module 2 (*in situ* H_2_O_2_ generator). Their combination creates a 3-input AND gate. In certain circumstances, candidate ‘bait’ proteins A, B, and C may not necessarily be totally distinct. (B-C) Assessment of split-APEX2 catalytic specificity by fluorescence imaging in 293T cells, with quantification of signal contrast between the stress granule (SG) interior and exterior. (D) Real-time imaging of light-activated PL in nucleoli. (E) Determination of the essential elements for light-activated catalysis (n = 40). (F) Real-time comparison between light-activated and H_2_O_2_-activated PL in live 293T cells. (G) Accumulation of catalytic contrast (Log_2_(△Contrast)) over time. (H) Western blot validation of the essential components for module 2. (I) Western blot comparing miniSOG and SOPP3 performance in module 2. (J) Western blot analysis of the performance of module 2 in the presence or absence of the SOD1 inhibitor LCS-1, confirming the dependence of module 2 on SOD1. Data in (C), (D), (G) are mean ± 95% CI (n > 50, n = 7, n > 11, respectively). Scale bars, 10 μm. GAPDH serves as a loading control in (H-J). Kruskal-Wallis test (C, E); ns, not significant, ***adj. p (BH corrected) < 0.001, ****adj. p < 0.0001, adj.p.

### A genetically encoded H_2_O_2_ generator enables in situ catalysis and improves specificity

Despite the improved specificity achieved with split-APEX2, this strategy still demands exogenous H_2_O_2_ supplementation, which can induce cytotoxic effects and provides limited spatiotemporal precision. To circumvent this, we developed an alternative two-input AND-gate system by coupling APEX2 with a light-controllable H_2_O_2_ generator, miniSOG-SOD1 (Figure 2A, Module 2).

We co-targeted APEX2 and the H_2_O_2_ generator in the nucleolus. As anticipated, blue light illumination triggered the progressive quenching of miniSOG fluorescence^41^—a characteristic indicator of superoxide anion production^42^—concomitant with the emergence of a catalytic signal detected by the membrane-permeable PL probe, Amplex UltraRed (converted into red resorufin upon peroxidase-catalyzed oxidization)^31^, specifically originating from within the nucleolar compartment (Figure 2D). The reaction kinetics were dependent on light intensity. Crucially, catalysis required the presence of both APEX2 and the H_2_O_2_ generator, confirming the formation of a two-input AND-gate (Figure 2E, H). Furthermore, the signal was diminished by the SOD1 inhibitor LCS-1^43^, supporting the proposed superoxide anion-to-H_2_O_2_ conversion mechanism (Figure 2J).

Comparing this optogenetic 2-input AND-gate with the conventional H_2_O_2_-based system, we found that light-controlled system produced a catalytic signal *in situ* within the nucleolus and exhibited accumulatively higher specificity at all time points (Figure 2F-G). In contrast, the signal in the H_2_O_2_-treated group originated in the cytoplasm and diffused inward, creating significant background (Figure 2F). Thus, the optogenetic AND-gate enhances catalytic specificity.

The choice of photosensitizer is important, as the efficacy of the proximity labeling reaction depends on both ROS generation efficiency and ROS species specificity. We compared miniSOG^44,45^ with its mutant SOPP3^41,45,46^, a more potent singlet oxygen generator used in the SOPP3-APEX2 system^33^. We hypothesized that singlet oxygen could directly react with biotin derivatives^47–49^, bypassing APEX2 and short-circuiting the AND-gate logic. Systematic comparison in nucleolar catalysis revealed that with biotin-phenol as the substrate, the SOPP3-mediated signal saturated quickly (∼60 s), whereas the miniSOG-mediated signal increased steadily over the 120 s time course, indicating more linear and controllable kinetics (Figure S2F-G). When the catalytic time was 2 minutes, the catalytic effects mediated by SOPP3 and miniSOG were nearly identical (Figure 2I). More importantly, when biotin-aniline was used as the substrate (Figure S2F-G), the “SOPP3 only” control group exhibited strong catalysis, nearly equivalent to the full SOPP3+APEX2 system. In contrast, the “miniSOG only” control showed low activity (Figure S2G). This confirms that SOPP3’s singlet oxygen directly catalyzes biotin-aniline labeling, compromising the integrity of the AND-gate. Given the utility of biotin-aniline and its derivatives for labeling targets like RNA^48,50^, this APEX2-independent activity makes SOPP3 unsuitable for true AND-gate constructions. Therefore, despite its slower catalytic rate, miniSOG is the superior choice for preserving the integrity of the logic gate.

### Phase-APEX2: a 3-input AND-gate for highly specific MLO labeling

To achieve maximal labeling specificity, we engineered Phase-APEX2, a 3-input AND-gate that combines our split-APEX2 (Module 1) and light-inducible H_2_O_2_ generator (Module 2). Imaging-based analyses in both nucleoli and stress granules revealed a clear specificity hierarchy: the 3-input gate significantly outperformed 2-input gates, which themselves were superior to single-bait strategies. Remarkably, each additional logical input conferred an approximate two-fold specificity enhancement (Figure 3A, S3A).

**Figure 3.**
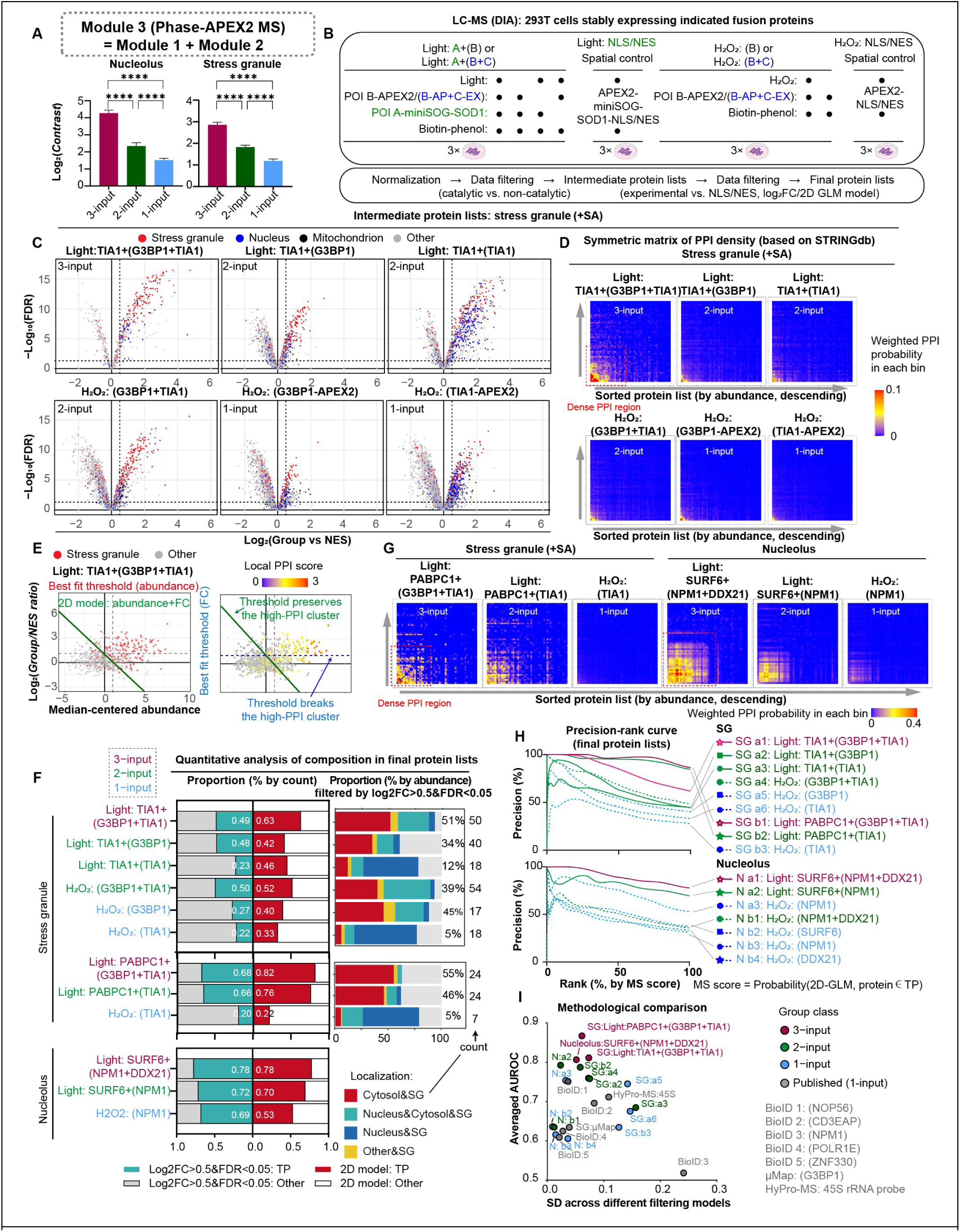
Phase-APEX2-MS and a 2D filtering approach improve specificity for MLO proteomics. (A) Catalytic contrast comparison for 1-, 2-, and 3-input AND gates. Data are mean ± 95% CI (n > 100); Kruskal-Wallis test; ****adj. p (BH corrected) < 0.0001. (B) Experimental design. Volcano plots of proteomes from different PL strategies. (D) Weighted joint probability heatmaps showing the distribution of TP (true-positive)-enriched, confidence(STRINGdb)-weighted protein-protein interactions across the abundance-rank space. Warmer colors indicate bins with higher probability. (E) Schematic showing different data filtering methods in the 2D scatter plot. Proteins with higher local PPI scores are corresponding to the dense interaction module (D). (F) Left: proportion of TP proteins from different PL strategies. Right: abundance distribution of SG subpopulations within the TP set. (G) Weighted joint probability heatmaps (as described in D, TP: SG/Nucleolus). (H) Precision-Rank curves. Overall specificity and robustness of PL strategies across different filtering models (1D log_2_FC, 1D abundance, and 2D log_2_FC+abundance, three models are shown in E).

To comprehensively quantify on-target specificity, we performed DIA-MS analysis in SGs and the nucleolus, two well-characterized MLOs with extensive proteomic data from orthogonal methods^26,36,37,51^. Following data normalization, we generated intermediate protein lists for each experimental group through data filtering based on catalytic versus non-catalytic conditions and examined the data distribution patterns across these groups (Figure 3B).

In our initial experiments (Batch 1), we evaluated 1-input, 2-input, and 3-input strategies using two canonical SG markers, G3BP1^7^ and TIA1^8^. Volcano plot analysis demonstrated that compared to 1-input and 2-input systems, the 3-input Phase-APEX2 system positioned more true-positive (TP) SG proteins closer to the upper-right quadrant with higher statistical significance, while simultaneously reducing contamination from off-target nuclear and mitochondrial proteins (Figure 3C). Further MS experiments (Batches 2 and 3) using three distinct markers for both SGs and nucleoli corroborated these findings, confirming that the 3-input strategy consistently achieves higher enrichment and statistical power for target proteins (Figure S3C).

MLOs are characterized by high molecular density^52^, which facilitates dense protein-protein interaction (PPI) networks. A highly specific labeling method should capture the dense network feature. To visualize the dense network, we generated PPI density heatmaps by annotating our proteomic data with the high-confidence PPI information from STRING database^53^, with proteins on both the x- and y-axes sorted by descending abundance. Strikingly, a large, high-density PPI hotspot was observed exclusively in the 3-input Phase-APEX2 datasets across all experimental batches, likely reflecting the high molecular density nature of MLOs (Figure 3D, 3G). In 2D scatter plots (abundance vs. log_2_FC), these high-density interactors co-localized with known MLO components in the upper-right quadrant (Figure 3E, S3E). This position (relatively high abundance and log_2_FC) aligns with the dense, phase-separated nature of MLOs.

To generate the final protein list preserving the high-density interaction region, we tried to filter out background proteins using a conventional threshold based on log_2_FC (experimental group vs. NLS/NES control). Unfortunately, we found that conventional data filtering based on an optimal log_2_FC threshold (AUROC analysis) often fragments the high-density interaction region (Figure 3E, S3E), implying it’s necessary to integrate both the enrichment factor and relative abundance in the analysis workflow.

To specifically address this challenge in MLO proximity-labeling proteomics, we developed a 2D General Linear Model (GLM), an unsupervised method that integrates both enrichment (log_2_FC) and relative abundance to define final protein lists. The 2D GLM more effectively preserved the integrity of the high-density interaction region while distinguishing true positives from background (Figure 3E, S3E). It also generates a probabilistic “MS score” for each protein. A direct comparison showed that our 2D GLM consistently outperformed single-threshold filtering, achieving higher sensitivity at constant specificity, and *vice versa* (Figure S3B), which underscores the importance of tailoring the analytical workflow to the high molecular density property of MLOs. It is worth noting that this specialized filtering method may not be necessary for conventional AP-MS experiments.

We employed different data filtering methods (2D GLM model or log_2_FC model) to generate final protein lists, which were then used to comprehensively compare the performance of the 3-input, 2-input and 1-input systems. With both filtering models, the 3-input Phase-APEX2 system was superior, but the specificity gain was most pronounced with the 2D GLM (Figure 3F). Interestingly, we found that constructing the logic gate with at least two distinct markers (G3BP1+TIA1) effectively mitigated the detection bias towards nuclear proteins introduced by TIA1 (H_2_O_2_:TIA1; Light:TIA1+(TIA1)).This finding was consistent with our imaging assessments (Figure S1D-F, S2A). Additionally, the 3-input system detected the highest number of SG components primarily localized to the cytoplasm (Cytosol & SG), indicating the lowest bias. Furthermore, because accuracy is sensitive to threshold selection, we used the more robust Precision-MS score curves and confirmed that the precision of the 3-input system remained higher than all other strategies across decreasing thresholds and surpassed previously published methods like BioID^36^, HyPro-MS^37^, and μMap^26^ (Figure 3H, S3D).

Finally, we assessed overall performance using Area Under the Receiver Operating Characteristic (AUROC) curves. The 3-input Phase-APEX2 system consistently achieved the highest average AUROC (0.8–0.9) with relatively low standard deviation (0–0.1), comprehensively demonstrating its superior specificity and robustness (Figure 3I).

### Phase-APEX2 maps the core interactome of MLOs

We hypothesized that the high-density interaction network identified by Phase-APEX2 represents the core structural and functional components of MLOs. To test this, we compared our proteomic data with other published MLO datasets. As mentioned above, our 2D GLM data filtering method can preserve the high-PPI cluster in 2D scatter plots, whereas traditional log_2_FC-based thresholding disrupts this high-PPI cluster (Figure 3E, S3E). Euler diagrams revealed that the Phase-APEX2 dataset, particularly when processed with our 2D GLM rather than the conventional threshold, consistently occupied a central, overlapping position among the datasets generated by all methods, strongly suggesting that it identifies the core MLO interactome (Figure 4A-B, Supplementary Table S1).

**Figure 4.**
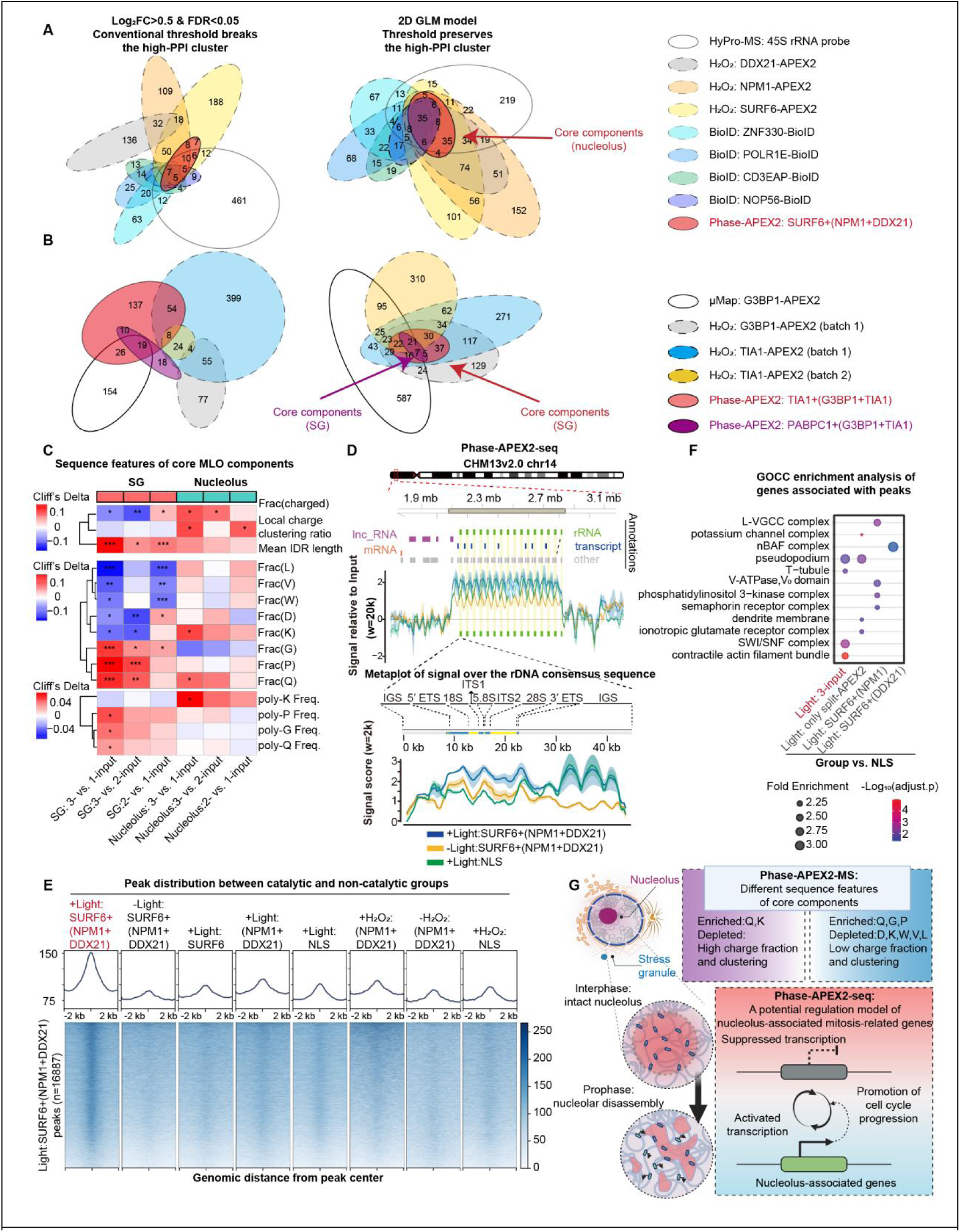
Phase-APEX2-MS/-seq identify core components of the nucleolus. (A-B) Euler diagrams of protein overlap from different PL datasets for the nucleolus (A) and SGs (B). (C) Sequence feature analysis of core MLO proteins. Cliff’s Delta is a non-parametric effect size quantifying the difference between two groups. Mann-Whitney U test; *adj.p (BH corrected) < 0.01, **adj. p < 0.001, ***adj.p < 0.0001. (D) Specificity of Phase-APEX2-seq. Top: signal enrichment on the rDNA array. Bottom: metaplot over the rDNA consensus sequence. (E) Signal distribution at peak regions. Top: metaplots of average signal intensity (±2 kb). Bottom: heatmap of signal intensity for all individual peaks (n = 16,887) in the 3-input group. (F) GOCC enrichment of peak-associated genes from experimental groups vs. NLS control. (G) Schematic showing features of SGs and nucleoli revealed by Phase-APEX2-MS and Phase-APEX2-seq.

### A distinct molecular grammar governs SG and nucleolar assembly

Given the significantly higher specificity of the Phase-APEX2 (3-input) system, we reasoned that a comparative analysis of protein sequences in different datasets could identify key MLO sequence grammars, which are likely obscured by the high background of off-target proteins in the 1- and 2-input analyses.

Sequence analysis (Figure 4C) revealed that the core SG proteins identified by our 3-input Phase-APEX2 method were enriched in glycine (G), proline (P), and glutamine (Q) but depleted of hydrophobic and charged residues. They also possessed longer intrinsically disordered regions (IDRs) and lacked charged blocks. In contrast, core nucleolar proteins were enriched in lysine (K) and displayed prominent charge block features. This highlights fundamental differences in the molecular grammar governing these two MLOs. Our finding aligns with parallel research in our lab on the immiscibility of different IDR-driven condensates^54^, and suggests that the molecular grammar of each MLO may be unique. Notably, recent studies based on IDR sequence analysis also identified K-patches as a unique sequence feature of nucleolar proteins^55^ and charge blocks as a primary characteristic of nuclear MLOs^55,56^, which is consistent with our findings.

By using the G/P/Q-rich sequence feature combined with a high 3-input MS score threshold (>0.6), we identified 30 co-enriched proteins that included many well-known key SG components and several candidate proteins (Figure S4A-C, Supplementary Table S2). Among these 30 key SG proteins with high G/P/Q features, several paralogs were present, such as YTHDF1/2/3, PRRC2A/B/C, PUM1/2^57,58^, TNRC6A/B^59^, G3BP1/2^6,7^, and ATXN2/ATXN2L^60,61^. Most of these proteins are known to be key components of SGs, including scaffold proteins like G3BP1/2^6,7^, key regulators like ATXN2/ATXN2L^60,61^, and YTHDF1/2/3^62^. Interestingly, proteins with significant G/P/Q amino acid features such as KHSRP, MEX3A, SF1, FAM120C, CPEB2, BAG4, TRIP6, and RBM33 have not been sufficiently studied for their relevance to SGs, making them promising candidates for further investigation. Notably, the amino acid features enriched in core SG proteins (G/P/Q enrichment with reduced D/K/W/V/L and non-significant R/Y/F) do not completely overlap with the features commonly enriched in RBPs^63–66^, especially those in the nucleolus (high charge fraction and clustering propensity). Furthermore, some studies have separately shown that G-rich^67,68^, P-rich^68,69^ or Q-rich^61,68,70^ regions play critical roles in LLPS. Therefore, this indicates that the G/P/Q enrichment is a specific feature of the core set of SG constituents, not a general characteristic of all RBPs. This suggests that the high specificity of Phase-APEX2 (3-input) is also highly beneficial for analyzing the molecular grammar of MLO components (Figure 4G).

### Phase-APEX2-seq for in situ mapping of MLO-associated DNA

Mapping the genomic loci that organize nuclear MLOs is challenging, as traditional methods like TSA-seq^71^ are incompatible with live cells, and suffer from fixation artifacts. To overcome this, we developed Phase-APEX2-seq to capture MLO-associated DNA regions in living cells. We identified biotin-naphthylamine^50,72^ as a superior substrate over biotin-phenol (Figure S4E) and confirmed DNA modification via DNase I sensitivity. To apply Phase-APEX2-seq to the nucleolus, we used a 90-second light activation in living cells, followed by rapid lysis, DNA fragmentation, and enrichment of biotinylated fragments for sequencing.

Our method successfully detected nucleolar-associated rDNA arrays with high specificity (Figure 4E, S4F). The signal was strongly enriched across rRNA transcription regions, while remaining at background levels in the less conserved intergenic spacers (IGS)^73^, comparable to the NLS control (Figure 4E). Analysis of enriched peaks confirmed that the photocatalytic signal was higher than all non-catalytic, spatial (NLS), and H_2_O_2_-catalyzed controls, highlighting the superior specificity of Phase-APEX2-seq (Figure 4F). A systematic comparison of 1-, 2-, and 3-input systems was consistent with our proteomics data: the 3-input system generated longer average peak lengths (∼1.6–1.8 kb) and labeled a relatively high proportion of rDNA regions, demonstrating maximal specificity (Figure S4G).

### Annotation of labeled genomic regions links nucleolar dynamics to cell cycle progression

Functional clustering of genes near the identified peaks revealed that the DNA regions labeled by the Phase-APEX2 (3-input) system were significantly enriched for two functional clusters: ’SWI/SNF complex’ and ’contractile actin filament bundle’, compared to the control groups and 2-input photocatalysis datasets (Figure 4F). Previous studies have shown that many genes related to cell division are transcriptionally upregulated during the G2/M phase^74–76^, which coincides with nucleolar disassembly. These published findings partially overlap with our identified genes (Supplementary Table S1), including *BCL7B*, *ACTB*, *SMARCD1*, *SMARCC1*, *SMARCD2* (SWI/SNF complex), *FBLIM1*, *PDLIM1/7*, *MYH9*, *MYH14*, *LDB3*, *CNN2*, *ACTN1*, *GAS2L1*, *TRIP6*, *TPM4* (contractile actin filament bundle), and *NDE1*, *RACGAP1*, *PPP1CC*, *GPSM2*, *SAPCD2*, *MELK*, *CEP55* (cell cycle-related). As it has been shown that a significant portion of Nucleolus-Associated Domains (NADs) detach from the nucleolus and become transcriptionally active upon loss of nucleolar integrity^77,78^, we hypothesize that the mitosis-related genes identified here may be transcriptionally activated concurrently with nucleolar disassembly, thereby driving cell cycle progression (Figure 4G).

We believe that the use of multiple nucleolar markers in Phase-APEX2 (3-input) enhances both specificity and stability, allowing for a more specific and comprehensive identification of MLO-associated DNA regions within the nucleolus, thereby facilitating the discovery of gene regions closely related to biological functions.

## Discussion

In this study, we demonstrated that conventional single-bait proximity labeling is inadequate for specific MLO analysis which may be due to the dynamic nature of MLOs, a lack of unique markers for different MLO types, and an unfavorable concentration-volume trade-off. To overcome this, we developed Phase-APEX2-MS, a 3-input AND-gate method that leverages the co-enrichment of three catalytic elements. By integrating a split-APEX2 system with a light-controlled H_2_O_2_ generator, we created a stringent 3-input AND-gate platform for rapid (30–120 s), light-inducible labeling that profoundly increases proteomic specificity in nucleoli and stress granules.

Furthermore, Phase-APEX2-MS’s high specificity enables the identification of protein components with high PPI propensity. To capture these components accurately, we introduced a 2D Generalized Linear Model (GLM) to the Phase-APEX2 workflow, which successfully identifies the core components of MLOs. Sequence analysis of these proteins revealed distinct sequence features between SG and nucleolar proteins.

We also extended the platform to Phase-APEX2-seq for direct DNA proximity labeling in living cells. This method enables rapid (90 s) photocatalysis with superior specificity over H_2_O_2_-based catalysis. Consistent with our proteomics data, the 3-input strategy offered the highest specificity for identifying nucleolus-associated DNA, including a cluster of mitosis-related genes upregulated during the G2/M phase. In conjunction with similar phenomena reported in previous studies^74–78^, this finding suggests a mechanistic link between nucleolar disassembly and the transcriptional activation of these genes during mitosis.

In summary, obtaining high-fidelity data remains a central challenge when characterizing dynamic systems like MLOs. The evolution of methodologies is clearly trending toward live-cell analysis with ever-increasing specificity. Phase-APEX2 represents a key advance on this trajectory, offering a powerful and versatile toolkit to unravel the composition and function of MLOs.

## Resource availability

Requests for further information and resources should be directed to and will be fulfilled by the lead contact, Pilong Li.

## Materials availability

Unique reagents and stable cell lines generated in this study are available from the lead contact.

## Data and code availability

- The mass spectrometry proteomics data reported in this paper have been deposited in the iProX database (https://www.iprox.cn/) of the National Proteome Science Center, Beijing, with the dataset identifier IPX0013699000.
- The raw sequencing data for Phase-APEX2-seq reported in this paper have been deposited in the Genome Sequence Archive (GSA, https://ngdc.cncb.ac.cn/gsa/) database under accession number PRJCA050735.

## Acknowledgement

We thank Peng Zou (College of Chemistry and Molecular Engineering, Peking University) and Ruijun Tian (Department of Chemistry, Southern University of Science and Technology) for generously providing the miniSOG plasmid and biotin-naphthylamine reagent (for the initial optimization of Phase-APEX2-seq protocol). We acknowledge the support of the Protein Chemistry and Proteomics Facility at Tsinghua University and the Proteomics Technological Platform of National Center for Protein Sciences (Beijing) for providing mass spectrometry services. This work was supported by grants from Natural Science Foundation of China (32125010, 32330024, 32450600 to P.L.; 92478128, 22477066 to W.Q.), and National Key R&D Program (2025YFA1308800, 2023YFF1204703 to P.L.; 2024YFA1308000 to W.Q.), and Natural Science Foundation of Beijing (Z230014 to P.L.; JQ25018 to W.Q.), and the Fundamental Research Funds from Beijing National Laboratory for Molecular Sciences (BNLMS202301 to W.Q.) and the Shenzhen Medical Research Fund (B2401004 to W.Q.).

## Author contributions

Z.C., W.Q., and P.L.conceived and designed the study and wrote the manuscript draft. Z.C. performed the majority of the experiments, including cell culture, plasmid construction, microscopy, immunoblotting, and sample preparation for MS/NGS experiments, and conducted data analysis. Z.C. and Y.C. optimized the initial proteomics workflow and performed the proteomics experiments for the nucleolus and stress granule systems (batches 2 and 3). Z.C., M.W., and B.F. performed the proteomics experiments for the stress granule system (batches 1 and 4). H.D. helped Z.C. to optimize protocols for cell culture and proximity labeling, and validated the performance of modules 1 and 2. Z.C. performed the proteomics data analysis. Z.C. optimized the sample preparation protocol for Phase-APEX2-seq, and M.H. performed the data analysis for Phase-APEX2-seq. W.Q., M.W., L.Z. provided the mass spectrometry analysis platform. W.C. provided some essential advice on experiment design. P.L. supervised all aspects of this study.

---

**Figure S1.**
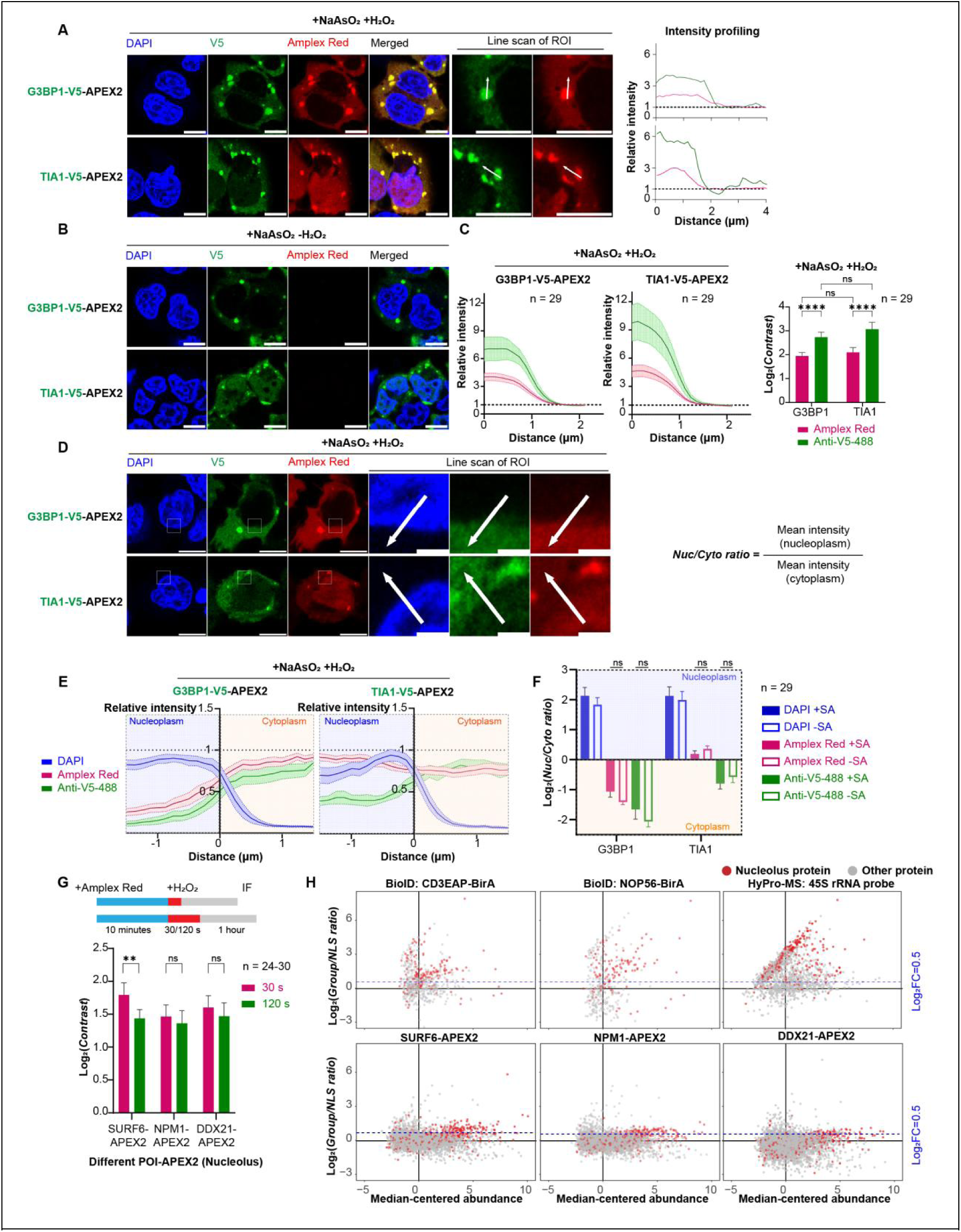
Challenges of applying conventional proximity labeling in stress granules and the nucleolus. (A) Immunofluorescence of POI-V5-APEX2 (POI: G3BP1/TIA1) and proximity labeling (PL) products in 293T cells treated with sodium arsenite (NaAsO_2_). (B) No-H_2_O_2_ control. (C) Fluorescence quantification along multiple axes (representative examples are shown in panel A) and signal contrast comparison. (D) Nucleocytoplasmic distribution of catalytic products in 293T cells. (E) Average nucleocytoplasmic distribution curves (representative examples are shown in panel D). (F) Nucleocytoplasmic distribution ratios from (E). (G) Assessment of catalytic specificity at different catalytic times. Mann-Whitney test; **adj. p < 0.01. (H) 2D scatter plots of different PL-derived proteomes. Data in (C, E, F) are mean ± 95% CI (n = 29); data in (G) are mean ± 95% CI (n = 24-30). Kruskal-Wallis test; ****adj. p (BH corrected) < 0.0001; ns, not significant. Scale bars, 10 μm.

**Figure S2.**
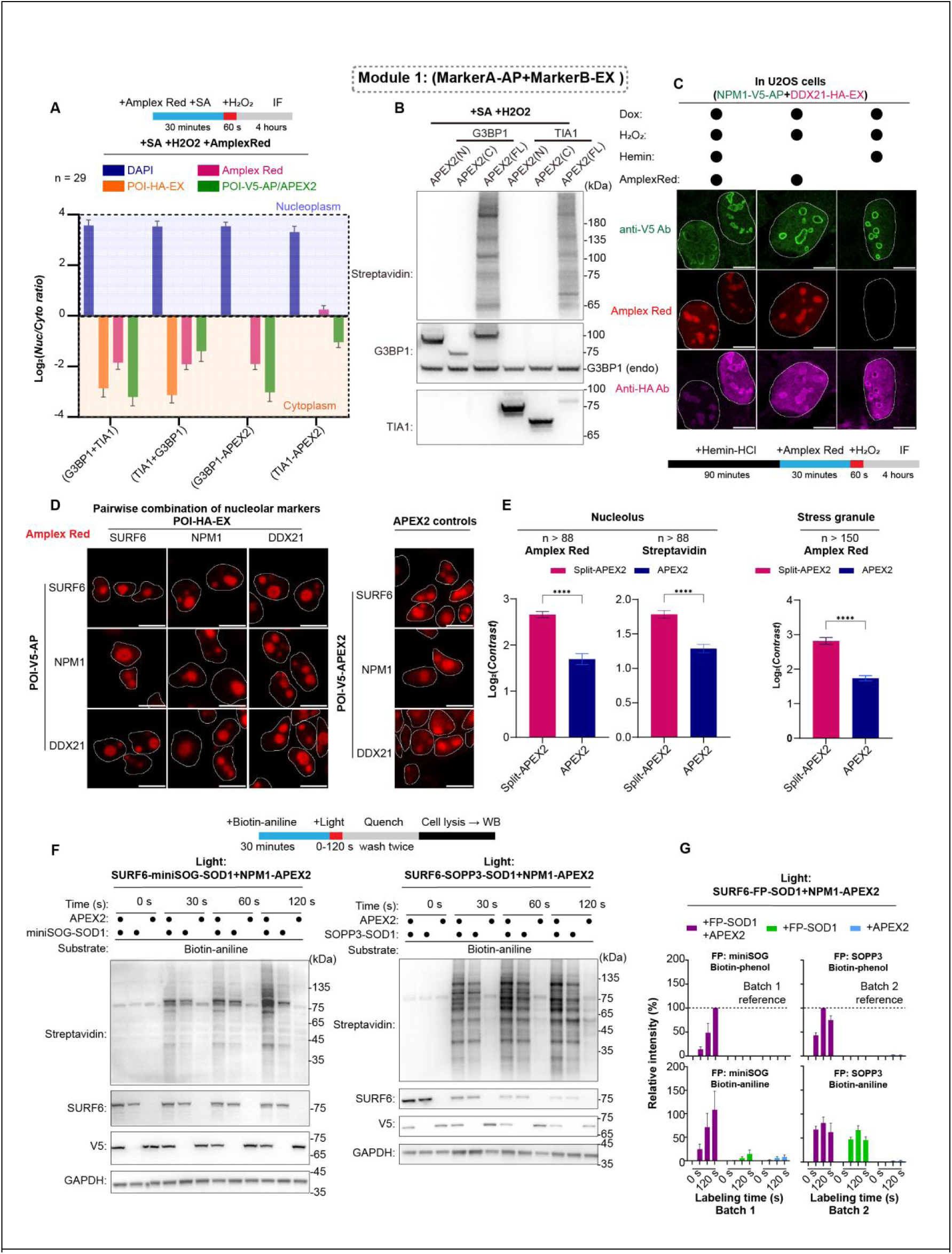
Characterization of split-APEX2 (Module 1) and the *in situ* H_2_O_2_ generator (Module 2). (A) Nucleocytoplasmic distribution ratios. (B) Western blot showing that individual split-APEX2 fragments are inactive. (C) Immunofluorescence showing that Module 1 activity is independent of exogenous hemin. (D) Catalytic specificity of split-APEX2 in the nucleolus (in 293T cells). (E) Quantification of catalytic specificity. Nucleolus, n > 88; stress granules, n > 150. Mann-Whitney test; ****p (BH corrected) < 0.0001. (F) Western blot evaluating the temporal response of Module 2 with different photosensitizers and substrates. (G) Protein dot blot comparing the performance of Module 2 with different photosensitizers and substrates. The maximum signal achieved with biotin-phenol in each group (miniSOG/SOPP3) was set to 100%. Data are mean ± SD (n = 3 biological replicates). Scale bar, 10 μm. G3BP1(endogenous) and GAPDH serves as the loading control in (B) and (F) respectively.

**Figure S3.**
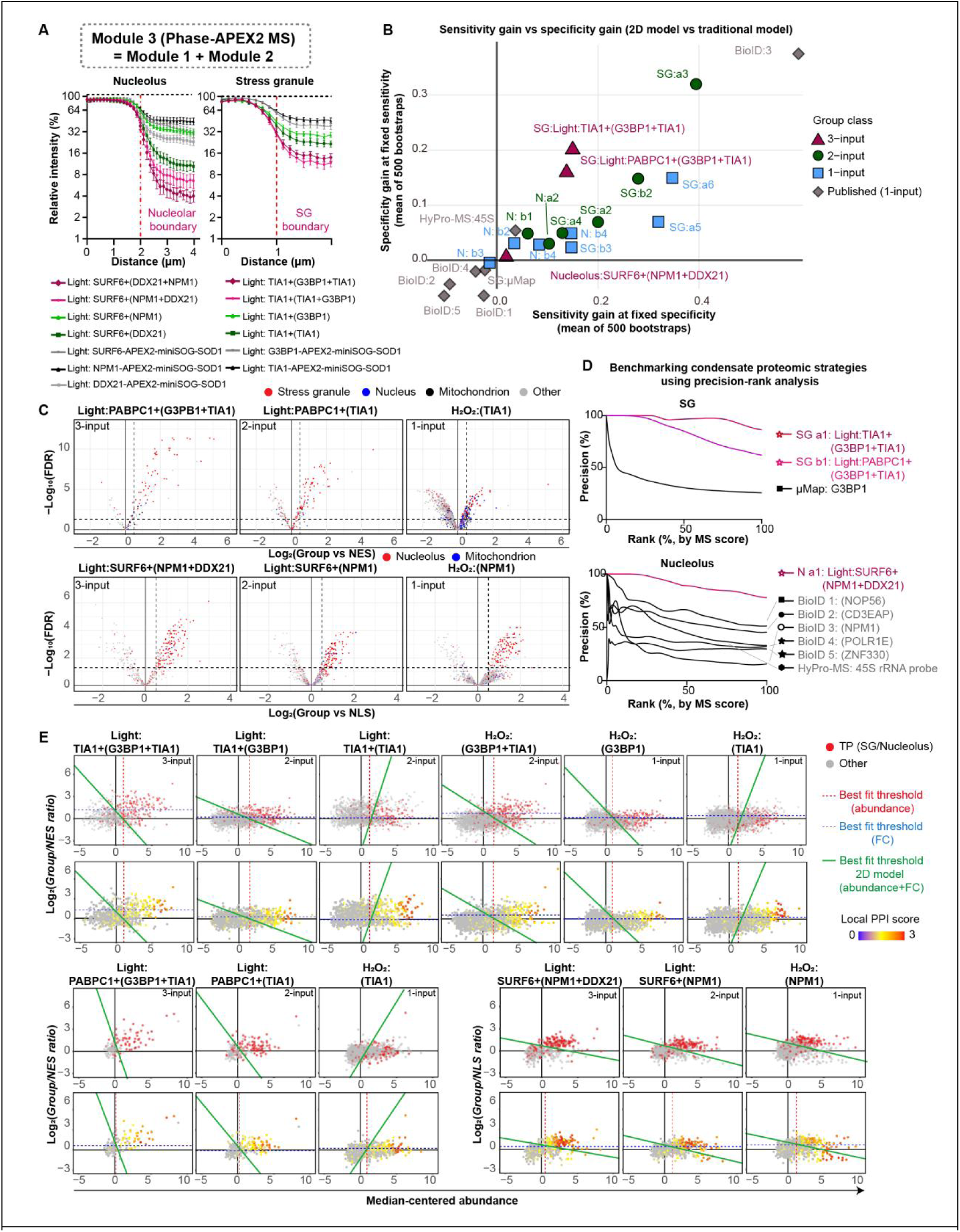
Performance evaluation of Phase-APEX2-MS in membraneless organelles. (A) Evaluation of catalytic specificity for module 3. Data are mean ± 95% CI (n > 50). (B) Benchmarking of the 2D GLM vs. a standard threshold (log_2_FC > 0.5 & FDR < 0.05). (C) Volcano plots of different PL-derived proteomes. (D) Precision-Rank analysis benchmarking different PL strategies. (E) Supplementary 2D scatter plots, analogous to Figure 3E.

**Figure S4.**
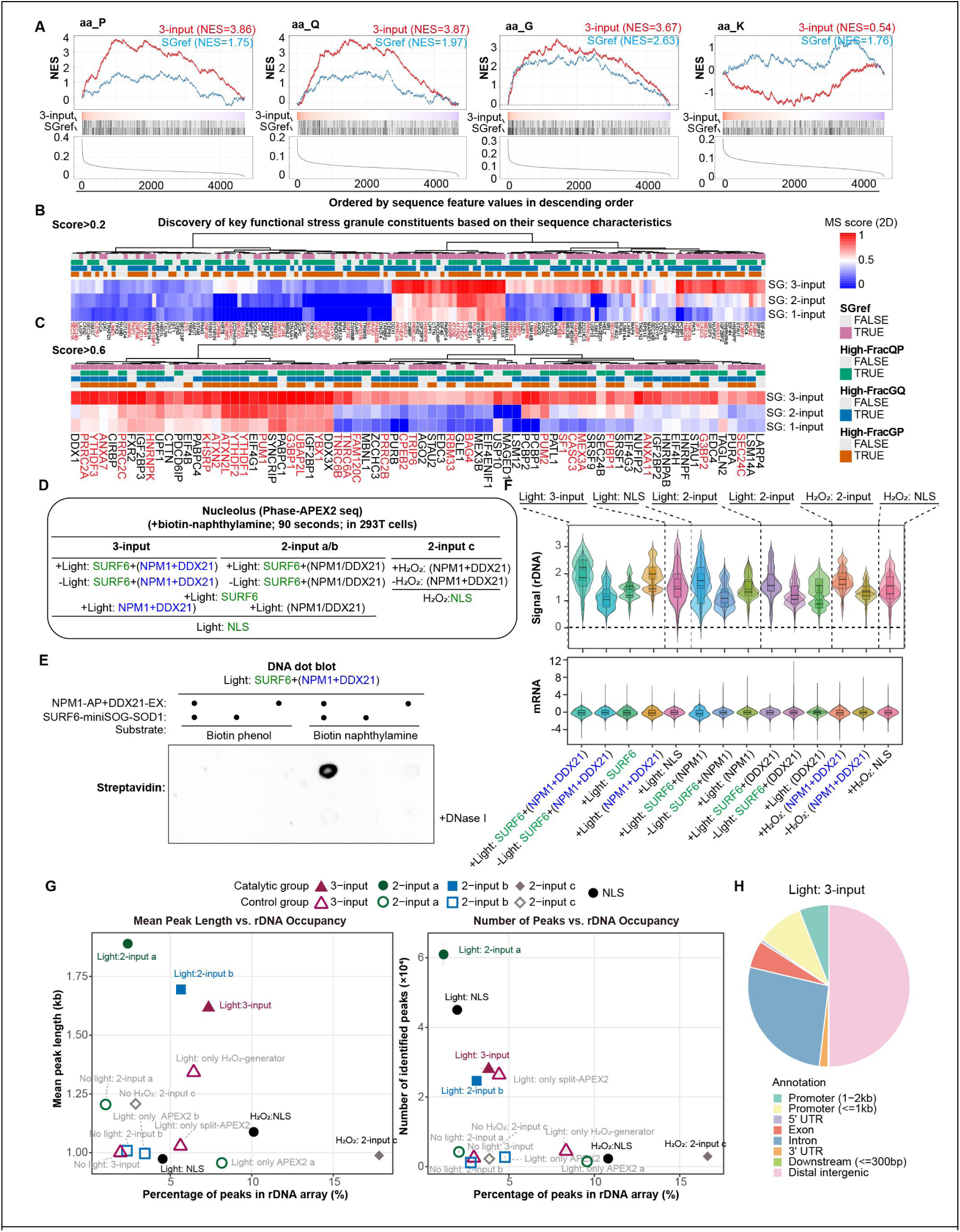
Supplementary data for sequence feature analysis of core SG proteins and Phase-APEX2-seq. (A) GSEA-like analysis of amino acid composition of core SG proteins. (B-C) Heatmap of score SG proteins from different PL strategies, clustered by MS scores and filtered at > 0.2 (B) and > 0.6 (C). (E) Schematic of Phase-APEX2-seq experimental groups. (E) Genomic DNA dot blot verifying the effectiveness of Phase-APEX2-seq . (F) Violin plots showing Phase-APEX2-seq signal distribution across different conditions. (G) Scatter plots comparing mean peak length and total peak number against the percentage of peaks within the rDNA array per condition. (I) Pie chart showing the genomic annotation of peaks from the 3-input system.

## Method details

### Cell culture

HEK293T and U2OS cells were cultured in high-glucose DMEM (Cytiva, SH30243.01) and McCoy’s 5A medium (Sigma, M9309), respectively, supplemented with 10% fetal bovine serum (VivaCell, C04001-500). Cells were maintained in a 37°C incubator with 5% (v/v) CO2. All cell lines were tested negative for mycoplasma contamination. The medium color was not allowed to turn yellow anytime, and cell confluence was kept below 90% to prevent the accumulation of reactive oxygen species (ROS) or contact inhibition that could interfere with subsequent experiments. For all experiments involving the Tet-On system, protein expression was induced by adding doxycycline hyclate (Solarbio, D8960) to a final concentration of 0.03-1 µg/mL (according to the expression efficiency of plasmids) 20 hours prior to the experiment. Once induced by adding doxycycline hyclate, cells were not cultured further to avoid potential accumulative stress responses.

### Contructs

Plasmids containing the full-length sequences of miniSOG and SOPP3 were gifts from the laboratories of Peng Zou and Wei Qin, respectively, and were used for PCR amplification and subsequent plasmid construction. Other gene sequences were either amplified by PCR from a high-quality cDNA library derived from HEK293T cells or obtained directly through gene synthesis (Tsingke, standard gene synthesis service). All sequences were verified by Sanger sequencing (Tsingke) focusing on the target region before plasmid construction. A modified Piggybac system plasmid (Addgene,122267) , containing the transposon-binding sequences flanking the main transcription region, was used as the backbone for all constructs. This backbone was optimized with a modified promoter to reduce leaky expression, and the original hygromycin resistance gene was replaced with Blasticidin and Puromycin resistance genes to facilitate drug selection for triple-plasmid co-expression systems. Target fragments were recombined with the backbone vector using a seamless cloning kit (Abclonal, RK21020).

After two rounds of selection, the abundance and integrity of expressed fusion proteins were verified by immunoblotting. Expression levels were adjusted to be comparable to or a bit higher than their endogenous counterparts. If catalytic activity was low, we re-checked protein integrity, replaced the fusion tag, or increased H_2_O_2_ generator expression. Following three rounds of selection, antibiotics were removed, and cells were cultured for an additional 72 hours before being cryopreserved or used in experiments. To prevent gene silencing and ensure reproducibility, all stable cell lines were used within two months of establishment.

### Generation of stable cell lines

Only stable cell lines were used for subsequent experiments.

To generate stable cell lines, cells at 80% confluence were co-transfected with the expression plasmid(s) and the Piggybac transposase (Pb) plasmid at a 3:1 mass ratio using Lipo8000 transfection reagent (Beyotime, C0533). For multi-bait systems, cell lines were generated sequentially, starting with the APEX2/AP/EX constructs, followed by the H_2_O_2_ generator constructs. A maximum of two expression plasmids were co-transfected with the Pb plasmid at one time. The total amount of plasmid DNA did not exceed 2 µg per well in a 6-well plate. The medium was replaced 12 hours post-transfection. After another 24 hours, cells were subcultured 1:3 and selected with Hygromycin B (200 µg/mL; MedChemExpress, HY-B0490), Puromycin (1 µg/mL; MedChemExpress, HY-B1743), or Blasticidin S HCl (15 µg/mL; Beyotime, ST018).

After two rounds of selection, the abundance and integrity of expressed fusion proteins were verified by immunoblotting. Expression levels were adjusted to be comparable to or a bit higher than their endogenous counterparts. If catalytic activity was low, we re-checked protein integrity, replaced the fusion tag, or increased H_2_O_2_ generator expression. Following three rounds of selection, antibiotics were removed, and cells were cultured for an additional 72 hours before being cryopreserved or used in experiments. To prevent gene silencing and ensure reproducibility, all stable cell lines were used within two months of establishment.

### Proximity labeling reaction conditions

The choice of catalytic substrate depended on the downstream application. For detecting protein labeling products, either 500 µM biotin-phenol (MedChemExpress, HY-125658) or 500 µM biotin-aniline (Iris-Biotech, LS-3970) was used. For detecting DNA labeling products, 500 µM biotin-naphthylamine (Iris-Biotech, LS-4650) and 500 µM biotin-phenol were used. For microscopy imaging, 500 µM biotin-phenol or 50 µM Amplex UltraRed (Invitrogen, A36006) was used. In the stress granule system, 1 mM sodium arsenite was added along with the biotinylated substrate to induce stress granule formation for 30 min.

In experiments validating split-APEX2 and conventional APEX2 (Module 1), cells were induced with 1 µg/mL doxycycline hyclate (Solarbio, D8960) for 20 hours to express the proximity labeling-related fusion proteins. The medium was then replaced with fresh medium containing the catalytic substrate, and cells were incubated at 37°C for 30 min (for biotin-phenol, biotin-aniline, biotin-naphthylamine) or 15 min (for Amplex UltraRed). H_2_O_2_ (Sigma, 323381) was then rapidly added to a final concentration of 1 mM and mixed as soon as possible. After 1 min, the reaction was quenched by washing the cells three times with Quenching Solution. The Quenching Solution contained 10 mM sodium ascorbate (Aladdin, S105026), 5 mM Trolox (MedChemExpress, HY-101445), and 10 mM KN3 (Macklin, P922726).

For photo-activated proximity labeling experiments, a 460-465 nm blue LED light panel (compatible with standard 96-well plates, with 96 uniformly distributed LED bulbs, 5-15 mW/cm2, Xuzhou Aijia Electronic Technology, Taobao) was used to initiate the reaction. After illumination, the reaction was immediately terminated by washing the cells three times with Quenching Solution.

### Immunofluorescence detection of proximity labeling products

After quenching the reaction, different immunofluorescence protocols were chosen based on the method for detecting biotinylated products.

For detection using fluorescently-conjugated streptavidin (IF-protocol A), the basic procedure was similar to published studies^79^ with minor modifications. Cells were fixed with 4% PFA (Leagene, DF0131) for 5 min, followed by rapid fixation and permeabilization with ice-cold methanol for 15 min. Cells were further permeabilized with PBS containing 0.5% (v/v) Triton X-100 for 10 min. After washing three times with PBS (Sangon, E607008), cells were blocked with 5% (w/v) BSA in PBST (PBS with 0.1% Tween-20) for 1h. Cells were then washed three times with PBST and incubated with primary antibody solution at 4°C for 16 h. The primary antibodies included: Mouse anti-V5 (Sino Biological, 100378-MM04, 1:500), Rabbit anti-V5 (Proteintech, 14440-1-AP, 1:500), Mouse anti-HA (Proteintech, 66006-2-Ig, 1:500), Streptavidin-YSFluorTM 488 (Yeasen, 35103ES60, 1:500), Streptavidin-YSFluorTM 594 (Yeasen, 35107ES60, 1:500), and Streptavidin-YSFluorTM 647 (Yeasen, 35104ES60, 1:500). After primary antibody incubation, cells were washed four times with PBST and incubated with secondary antibody solution for 4 h. The secondary antibodies included: Goat anti-Rabbit-IgG H&L (Alexa Fluor® 488) (Proteintech, RGAR002, 1:500), Goat anti-Rabbit-IgG H&L (Alexa Fluor® 594) (Proteintech, RGAR004, 1:500), Goat anti-Rabbit-IgG H&L (Alexa Fluor® 647) (Proteintech, RGAR005, 1:500), Goat anti-Mouse-IgG H&L (Alexa Fluor® 488) (Proteintech, RGAM002, 1:500), Goat anti-Mouse-IgG H&L (Alexa Fluor® 594) (Proteintech, RGAM004, 1:500), and Goat anti-Mouse-IgG H&L (Alexa Fluor® 647) (Proteintech, RGAM005, 1:500). Cells were then washed three times with PBST, stained with DAPI solution (Solarbio, C0065) for 10 min, and finally washed three times with PBST to remove unbound dyes and secondary antibodies.

For visualizing the reaction location using Amplex UltraRed (IF-protocol B), the procedure was identical to published studies. Cells were fixed with 4% PFA for 10 min, washed three times with PBST, and permeabilized with 0.5% Triton X-100 in PBS for 15 min. After washing three times with PBST, cells were incubated with primary antibody solution for 1 h (same antibodies as in protocol A, but without streptavidin, as Amplex UltraRed produces a red fluorescent signal). After primary antibody incubation, cells were washed four times with PBST and incubated with secondary antibody solution for 1 h. Subsequent steps were the same as in Protocol A. If no primary antibody was used and only the Amplex UltraRed signal distribution was assessed, cells were directly stained with DAPI after permeabilization, washed three times with PBST, and imaged immediately on a Nikon A1HD25 confocal microscope using a 100× oil-immersion lens.

### Live-cell imaging of blue light-activated proximity labeling

Live-cell imaging was performed on a Nikon A1HD25 confocal microscope equipped with a live-cell incubation chamber (37°C, 5% CO2). The H_2_O_2_ generator was excited using a 488 nm laser at 5-20% of its maximum power. Signals were detected in line-by-line mode (Line 4-1) for three fluorescent channels (DAPI, miniSOG, Amplex UltraRed). The total exposure time was 5.8 s, with one frame captured every 6 seconds. No PMT High Voltage Offset was applied to any channel to avoid filtering out low fluorescence signals, which could lead to misinterpretation.

### Immunoblot analysis of proximity labeling

After quenching the reaction, cells were pelleted by centrifugation at 600 g for 5 min. The supernatant was discarded, and cells were lysed in RIPA buffer (50 mM Tris pH 8.0, 150 mM NaCl, 0.1% SDS, 0.5% sodium deoxycholate, 1% Triton X-100, 1× protease inhibitor cocktail [Selleck, B14001]). UltraNuclease (Yeasen, 20156ES50) was added to the RIPA lysate at a 1:200 (v/v) ratio, and the mixture was incubated at 37°C for 5 min to simultaneously lyse cells and digest RNA and DNA, which is helpful of releasing nucleic-acid associated proteins. Four volumes of acetone were added, and the mixture was inverted to mix and stored at -20°C for 16 h to precipitate and fully denature the proteins. The protein pellet was collected by centrifugation at 9000 g for 10 min and washed three times with 80% (v/v) acetone, ensuring complete removal of the supernatant each time to eliminate small reducing molecules (trolox, sodium ascorbate or free biotinlyated substrates)that could interfere with subsequent protein quantification. The remaining acetone was evaporated, and the protein pellet was completely dissolved in a high-concentration SDS and sodium deoxycholate RIPA solution (1% SDS, 2.5% sodium deoxycholate) with the aid of sonication (30 s ON/ 30 s OFF, 10 cycles). Finally, the sample was diluted tenfold with a dilution buffer (50 mM Tris pH 8.0, 150 mM NaCl, 0.25% sodium deoxycholate, 1% Triton X-100, 1× protease inhibitor cocktail).

All Western blot experiments were performed using 4-20% gradient precast gels (GenScript, M00655) for SDS-PAGE. Sample concentrations were normalized after quantification with a BCA assay (Yeasen, 20201ES76). After SDS-PAGE, proteins were transferred to a 0.2 μm PVDF membrane (Merck, ISEQ00010). The membrane was blocked with 5% (w/v) BSA in PBST at 4°C for 16 h, then washed four times for 5 min each with PBST to remove unbound BSA. The membrane was incubated with primary antibody in PBST (0.2% v/v Tween-20) on a shaker at room temperature. Primary antibodies included: Streptavidin-HRP (Proteintech, SA00001-0, 1:2000), Rabbit anti-V5 (Proteintech, 14440-1-AP, 1:1000), Rabbit anti-SURF6 (Sangon, D123258, 1:1000), Rabbit anti-GAPDH (Proteintech, 10494-1-AP, 1:1000), Mouse anti-MYC (Proteintech, 60003-2-Ig, 1:1000), Rabbit anti-G3BP1 (Proteintech, 13057-2-AP, 1:1000), and Rabbit anti-TIA1 (Proteintech, 12133-2-AP, 1:1000). After a 2 h incubation, the membrane was washed four times with PBST to remove residual primary antibody, followed by incubation with secondary antibody for 1 h on a shaker at room temperature. Finally, the membrane was washed five times for 5 min each with PBST and developed using an ECL reagent (ServiceBio, G2014, G2074, G2020). Secondary antibodies included HRP-conjugated Goat anti-Rabbit IgG (Proteintech, SA00001-2, 1:5000) and HRP-conjugated Goat anti-Mouse IgG (Proteintech, SA00001-1, 1:5000).

### Dot blot analysis of proteins and DNA

Protein samples were prepared as described for the immunoblotting experiments. DNA samples were prepared as described for the Phase-APEX2-seq experiments.

Protein samples, typically at a concentration greater than 500 ng/µL after quantification, were directly spotted (2.5 µL) onto a 0.2 µm nitrocellulose membrane (Amersham, 10600001) and allowed to air dry. DNA samples were adjusted to a concentration of 1 µg/µL and spotted (2 µL) onto a nitrocellulose membrane. The membrane was then cross-linked for 30 minutes under a UV lamp in a clean bench; the duration may need adjustment depending on the UV lamp’s power and position. The nitrocellulose membrane was blocked with 0.5% (v/v) BSA in PBST on a shaker at 4°C for 16 h, followed by a further blocking step with QuickBlock (Beyotime, P0222) on a shaker at room temperature for 1 h. The membrane was washed four times with PBST containing 0.2% (v/v) Tween-20 for 5 min each. It was then incubated with the primary antibody solution (Streptavidin-HRP in 0.2% Tween-20 PBST) on a shaker at room temperature for 1 h to allow antigen-antibody binding. The membrane was washed five times with 0.2% PBST for 5 min each time. Subsequent detection and developing steps were the same as in the immunoblotting experiments.

### Hemin dependence assay

After inducing fusion protein expression with doxycycline, Hemin-HCl (Sigma, 51280) was added to the culture medium to a final concentration of 5 µM (Hemin-HCl stock solution was prepared as previously described^31^). After 90 min, cells were carefully washed twice with fresh medium. Then, either 500 µM biotin-phenol or 50 µM Amplex UltraRed was added, and the cells were incubated for 15 min (strictly protected from light). Finally, the proximity labeling reaction was performed.

### Phase-APEX2-MS sample preparation

Due to the large number of samples, the experiments were conducted in batches. Each membraneless organelle system was processed in two batches, with each batch having a different research objective but including complete catalytic, non-catalytic, and spatial control groups, each with three biological replicates. For the stress granule system: Batch 1 (samples labeled SG a1-a6, using G3BP1 and TIA1 as baits) was prepared and analyzed using Protocol A; Batch 2 (samples labeled SG b1-b3, using PABPC1, G3BP1, and TIA1 as baits) was prepared and analyzed using Protocol B. For the nucleolus system: Batch 3 (samples labeled N a1-a3) used Protocol B; Batch 4 (samples labeled N b1-b4, corresponding to the APEX2-MS part in Figure 1D/E and Figure S1H) used Protocol A.

For all mass spectrometry batches, the protein extraction procedure was the same as that used for immunoblotting. After BCA quantification, the concentration of each sample was adjusted to 1 mg/mL, and 1 mg of protein was used for the subsequent streptavidin pull-down.

For each sample, 200 µL of streptavidin magnetic beads (ChomiX Biotech, 2030002) were used for pull-down enrichment. The beads were first washed twice with 1 mL of RIPA buffer. The sample solution was then added, and the mixture was incubated on a rotary shaker at 4°C for 16 h.

After enrichment, the beads were pelleted using a magnetic rack. The beads were washed twice with 1 mL of RIPA buffer, once with 1 mL of 1 M KCl, once with 1 mL of 0.1 M Na2CO3, once with 1 mL of 2 M urea in 10 mM Tris-HCl (pH 8.0), and twice with RIPA buffer. Finally, the beads were washed three times with 1 mL of 50 mM ammonium bicarbonate (ABC) solution.

The beads were resuspended in 200 µL of 6 M urea in 50 mM ABC buffer and subjected to the following steps on a thermomixer. 25 µL of 100 mM dithiothreitol (DTT, Yuanye, S11080) was added to each sample and incubated at 35°C, 1150 rpm for 30 min. The proteins were then alkylated by adding 25 µL of 200 mM iodoacetamide (IAA, Vetec, V900335) and incubating at 25°C, 1150 rpm for 30 min in the dark. The beads were then washed three times with 50 mM ABC and resuspended in 200 µL of 50 mM ABC buffer (containing 1 M Urea). For digestion, 4 µL (Protocol A) or 2 µL (Protocol B) of 1 µg/µL trypsin (Sigma, T1426) were added to each sample, and incubation was carried out on a thermomixer at 37°C, 1150 rpm for 4 h (Protocol A) or 16 h (Protocol B) overnight.

The eluted peptides were desalted on C18 StageTips (Waters, cat. no. WAT036905). The desalting columns were activated sequentially with methanol, acetonitrile, and 50% acetonitrile. After digestion, the beads were pelleted using a magnetic rack, and the supernatant was loaded onto the column. Before elution, the columns were washed 10 times with 0.1% formic acid in water. The peptides were then eluted sequentially with 300 µL of 50% acetonitrile, 80% acetonitrile, and 50% acetonitrile. The final eluate was dried in a vacuum concentrator and stored at -80°C. The protein sample was redissolved in 0.1% formic acid in water for LC-MS/MS data-independent acquisition (DIA).

#### Protocol A

A Dionex UltiMate 3000 Rapid Separation LC (RSLC) system (Thermo Fisher Scientific) was coupled to a timsTOF Pro2 mass spectrometer with a CaptiveSpray nano-electrospray ion source (Bruker Daltonics). A 100 ng peptide mixture was separated on integrated spray tip columns (75 μm i.d. × 25 cm) packed with 1.9 μm/120 Å ReproSil-Pur C18 resins (Dr. Maisch GmbH, Germany). The column was heated to 60°C. The flow rate was 300 nL/min over a 90-min gradient, except for the first 25 min, where it was 500 nL/min. Mobile phase A was water with 0.1% formic acid (FA), and B was 80%/20% acetonitrile/water with 0.1% FA. The gradient was: 0% B for 25 min, 6%–40% B for 60 min, 99.0% B for 3 min, and 0% B for 2 min. Samples were acquired in diaPASEF mode with the following settings: precursors were scanned from m/z 100-1700, with precursors between m/z 400 and 1200 isolated in 32 windows. The isolation window was 25 Th with 1 Da overlap. The ion mobility range was 0.6 to 1.6 Vs cm-2, and both accumulation and ramp times were 100 ms. The estimated cycle time was 1.8 s.

The DIA raw files were analyzed by Spectronaut™ (version 19.5) using directDIA mode against the human UniProt database (updated July 30, 2024; 20,436 protein groups) on a server with an AMD Ryzen Threadripper 3990X CPU and 1024 Gb RAM. Trypsin was the enzyme, with a maximum of 2 missed cleavages. iRT-RT regression was set to ‘deep learning-assisted iRT regression’. Precursor and protein q-value cutoffs were both set to 1%.

#### Protocol B

The procedure was identical to that reported for the BRET-ID method^80^. Peptides were separated using a loading column (100 µm × 2 cm) and a C18 separating capillary column (100 µm × 15 cm) packed in-house with Luna 3 µm C18(2) bulk packing material (Phenomenex, USA). The mobile phases (A: water with 0.1% formic acid; B: 94% acetonitrile with 0.1% formic acid) were driven by a Vanquish Neo UHPLC system. The LC gradient was held at 4% B for the first 4 minutes, followed by an increase from 5% to 20% B from 4 to 109 minutes, an increase from 20% to 35% B from 109 to 150 minutes, and an increase from 35% to 99% B from 150 to 159 minutes. For samples analyzed on Q Exactive-plus series Orbitrap mass spectrometers (Thermo Fisher Scientific), precursors were ionized using an EASY-Spray ionization source (Thermo Fisher Scientific) held at +2.0 kV, and the inlet capillary temperature was 320°C. Survey scans of peptide precursors were collected in the Orbitrap from 350-1800 m/z with an AGC target of 3,000,000, a maximum injection time of 20 ms, and a resolution of 70,000. For DIA, the resolution was set to 17,500. The AGC target for fragment spectra was 1,000,000 with an auto injection time. Normalized collision energy (NCE) was set at 28%. The default charge state was 3, and the fixed first mass was 200 m/z.

The raw data were processed using DIA-NN in an advanced library-free module. Parameters were mostly identical to BRET-ID^80^, except the MBR parameter and cross-run normalization was set to OFF.

### Phase-APEX2-seq experimental procedure

The genomic DNA extraction protocol was modified from the traditional phenol-chloroform method by using DNAzol to assist in cell lysis, which yields high-quality and high-concentration genomic DNA.

HEK293T cells were cultured in 10 cm dishes, with each dish representing one biological replicate. After the proximity labeling reaction, the reaction was rapidly terminated by washing the cells three times with 2 mL of Quenching Solution. Cells were collected using a cell scraper, centrifuged at 1000 g for 5 min, and the supernatant was discarded. Each sample was rapidly lysed with 2 mL of DNAzol reagent (Simgen, 3107100), and lysis was assisted by repeatedly pipetting with a needle until no visible clumps remained. Crude genomic DNA was extracted according to the DNAzol instructions, with care taken to remove as much supernatant as possible during centrifugation. The DNA was dissolved in 100 µL of 8 mM NaOH at room temperature for 1 h; the pellet did not dissolve completely at this stage and required multiple rounds of pipetting until the clump expanded into a cloud-like form (indicating protein contamination). The solution was diluted with 500 µL of low-EDTA TE buffer (Sangon, B548408), and 10 µg of RNase A (CWBiotech, CW0601S) was added to digest residual RNA at 37°C for 4 h in a Thermomixer (950 rpm). Then, 20 µg of Proteinase K (Sangon, B600169) was added to digest residual proteins at 55°C for 1 h in a Thermomixer (950 rpm). Finally, the solution was repeatedly extracted 3-5 times with DNA-grade phenol-chloroform reagent (Acmec, AC13309). During each extraction, the aqueous phase was carefully collected, avoiding the intermediate layer. If the intermediate layer was easily disturbed, it indicated significant protein contamination or high nucleic acid concentration, requiring an increase in volume to lower the concentration. To the resulting aqueous phase, one-tenth volume of 3 M NaOAc (Beyotime, ST342) and two volumes of ethanol were added. The mixture was incubated at -80°C for 16 h to precipitate the DNA. The DNA precipitate, visible as fibrous strands after inverting the tube 10 times, was pelleted by centrifugation at 13,000 g for 10 min to form a dense white pellet. The supernatant was removed, and the pellet was washed three times with 70% ethanol. After removing residual ethanol and air-drying, the DNA was dissolved in Low-EDTA TE with gentle pipetting (high-purity DNA dissolves completely within 1-5 min). The purity of the DNA solution was assessed: the 260/230 absorbance ratio was around 2.4-2.5, and the 260/280 ratio was around 1.8-1.85. Agarose gel electrophoresis showed almost no low-molecular-weight bands below 10 kb or smeared degradation bands, indicating high integrity and purity of the genomic DNA (data not shown). If the DNA was impure, it was re-digested with RNase A and Proteinase K, followed by another phenol-chloroform purification.

The quality of catalysis was assessed by dot blot analysis of the purified DNA. If catalysis was successful, subsequent steps were performed.

Purified genomic DNA was randomly fragmented using an enzymatic method (FS Module, Abclonal, RK20276). DNA solution and EP tubes were pre-chilled on ice. To 50 µg of DNA, 8 µL of FS Pro buffer I was added and mixed thoroughly. Then, 12 µL of pre-chilled FS Pro Enzymes II was rapidly added on ice, and the total volume was brought to 80 µL with Low-EDTA TE. The reaction was incubated at 32°C for 60 min. The reaction was stopped by adding EDTA to a final concentration of 50 mM. A 1 µL aliquot was taken to assess the fragment size distribution by agarose gel electrophoresis; DNA fragments should range between 200 bp and 1000 bp. Each sample was processed in two 80 µL reaction volumes, containing a total of 100 µg of genomic DNA. After stopping the reaction with EDTA, the sample was diluted to a total volume of 800 µL with TE buffer. 40 µL of Streptavidin magnetic beads (ChomiX Biotech, 2030002) were added, and the mixture was incubated on a rotary shaker at 4°C for 1 h to capture biotinylated DNA. The beads were then washed five times with B/W/T buffer (10 mM Tris-HCl, pH 8.0, 1 M NaCl, 0.1% (v/v) Tween-20). Finally, the beads were resuspended in 500 µL of Low-EDTA TE, and SDS (final concentration 0.2% v/v) and 40 µg of Proteinase K were added. The mixture was incubated at 65°C for 1 h to digest proteins. The DNA was then extracted twice with phenol-chloroform to facilitate its release from the magnetic beads, yielding an aqueous phase containing the free DNA. One-tenth volume of 3 M NaOAc (pH 5.2), 2.5 volumes of absolute ethanol, and 20 µg of glycogen (Beyotime, D0812) were added. The mixture was mixed well and incubated at -80°C for 48 h to precipitate the DNA. After centrifugation, the pellet was washed three times with 75% ethanol, air-dried, and dissolved in 20 µL of RNase/DNase-free H2O. For each sample, 1-10 ng of DNA was typically obtained for next-generation sequencing library preparation. Control groups with low catalytic activity yielded around 1 ng or less of DNA, which might be due to non-specific binding of residual nucleic acids to the magnetic beads or EP tube walls.

Library construction and sequencing services were provided by AZENTA, following a standard ChIP-seq library preparation protocol. Sequencing was performed on a NovaSeq 2×150 bp platform, generating approximately 8-10 Gb of data per sample.

Sonication was attempted for nucleic acid fragmentation, but the resulting DNA distribution was uneven, and the pull-down products had a much larger average molecular weight than the input. This was possibly due to biotinylation affecting the DNA double helix structure, making it unsuitable for library construction.

### Microscopy image analysis

All microscopy images were acquired using a Nikon A1HD25 microscope with a measurement depth of 12 bits. No PMT High Voltage Offset was applied to any fluorescence channel to avoid filtering out low fluorescence values, which could lead to misinterpretation.

Image analysis was performed using Fiji/ImageJ. For large-scale intensity profiling along a line, a custom macro script (Plot_MultiColor_TimeSeriesAnalysis.ijm) was used for automated measurement. The line width for measurement was 2-4 pixels. For each measurement, the mean of the 10 highest or lowest fluorescence values was used as a reference to normalize the results (reference value = 1).

### Quantitative analysis of biotinylated protein dot blots

Image analysis was performed using Fiji/ImageJ. For protein dot blot results, the sample spot position was determined based on the anti-GAPDH immunosignal, and the sample loading amount was measured. The Streptavidin-HRP signal was then measured, divided by the protein loading amount, and further divided by the highest signal value within the group to obtain the normalized dot blot signal. Each dot blot statistical result was based on three biological replicates performed on three different nitrocellulose membranes.

### Mass spectrometry data analysis

#### Public data used for analysis

For the µMap method, the median-normalized raw data from the supplementary materials^26^ of the original paper were used, and the normalization step was skipped in subsequent analyses. For the publicly available raw data from BioID-MS^36^ and HyPro-MS^37^, FragPipe software was used for protein identification and label-free quantification (LFQ), following the standard DDA-LFQ workflow. No Match-Between-Runs (MBR) or pre-normalization parameters were set, as the FragPipe documentation^81,82^ indicates that MBR is only suitable for intra-group correction, not for inter-group correction in proximity labeling AP-MS experiments.

#### Data annotation, normalization, and missing value imputation

First, the raw protein abundance matrix was read, duplicate protein names were removed, and the subcellular localization of each protein was annotated. Subcellular localization information from the Human Protein Atlas (HPA) database (v24.0)^16^ was used. Rows with “Uncertain” in the “Reliability (IF)” field were removed, and the “subcellular location” and “subcellular main location” fields were used for annotation. “Subcellular main location” was primarily used, while “subcellular location” was used to exclude secondary localizations. Mitochondrial localization was annotated using the MitoCarta3.0 database^83^. Stress granule localization was annotated using the “known SG reference” column from the “SG_ref” table in Supplementary Data 1 of the µMap paper, corroborated with the GO database^84^ entry GO:0010494 (cytoplasmic stress granule). The annotation priority was SG > nuclear > cytosol > nuclear_cytosol > mitochondrion > other (details in the original code). Sub-SG protein localization was annotated based on a combination of HPA_main_location and SGref (from the µMap paper supplement). Briefly, an SGref-annotated SG protein was named “cytosol&SG” if its HPA_main_location was cytoplasm, and so on.

After annotating subcellular localizations, the raw LFQ values were log2-transformed (ignoring null values, resulting in dataframe1). Quantile normalization was then performed within the three biological replicates of each group to reduce technical variation while preserving biological differences between groups. The data matrix was re-merged by group. Catalytic groups (e.g., photo-catalysis and H_2_O_2_ groups) and control groups (-light/-miniSOG-SOD1/-APEX2/-H2O2/NLS/NES) were defined. Missing values were imputed only in the control groups using a Perseus-type method^26^ for each column: missing values were replaced by random numbers drawn from a normal distribution down-shifted by 1.8 standard deviations with a width of 0.3 of each sample, resulting in dataframe2.

#### Inter-group differential analysis and initial ROC analysis (catalytic vs. non-catalytic control groups)

The differential expression between catalytic and control groups (excluding NLS/NES controls) was analyzed using the limma package to fit a linear model and apply empirical Bayes moderation. FDR values were adjusted using the Benjamini-Hochberg method. ROC curves were constructed using the pROC package^85^ based on predefined true positive (TP) and false positive (FP) sets to determine the optimal log_2_FC threshold. Significantly enriched proteins were filtered based on the ROC-determined log_2_FC threshold and an FDR < 0.05 (requiring at least two valid values in the three biological replicates of the catalytic group), resulting in dataframe3.

For the stress granule system, GO-annotated SG proteins were used as the TP set, and mitochondrial matrix proteins from the MitoCarta database were used as the FP set. For the nucleolus system, proteins annotated to the nucleolus in the “subcellular main location” field of the HPA database were used as the TP set, and mitochondrial proteins from the MitoCarta database were used as the FP set.

#### Comparison with NLS/NES groups and 1D/2D model construction

Global quantile normalization was performed on all groups in dataframe1. For each group, only the proteins that passed the filtering threshold in dataframe3 were retained, using their corresponding values from dataframe1. These were re-merged into a new data frame (dataframe4) for subsequent analysis.

Log_2_FC (vs. NLS/NES control) and FDR (BH-adjusted) were calculated using the limma package^86^. The final protein lists were obtained by further filtering using either traditional thresholds (log_2_FC > 0.5, FDR < 0.05) or by constructing 1D/2D models. For the 1D model, log_2_FC (vs. NLS/NES control) or abundance (mean-median transformed relative abundance) was used as the model variable, and the glm function from the pROC package was used to train a classification model to distinguish TP from others. For the 2D model, both log_2_FC (vs. NLS/NES control) and abundance were used as model variables to train a classification model using the pROC glm function. The model assigned each protein a probability score (MS score, 0-1) of belonging to the TP set and provided an optimal filtering threshold. The AUROC values and DeLong’s test were used to evaluate and compare the AUROC values of different methods and proximity labeling techniques (p-values were BH-adjusted). For the stress granule system, the TP set for training the 1D/2D models consisted of “Cytosol&SGs”, “Nuclear&Cytosol&SGs”, and “SGs&Other”. For the nucleolus system, the TP set consisted of proteins with “Nucleolus” in their HPA_location annotation.

#### PPI interaction density plot

PPI interaction heatmaps for different groups in dataframe4 were annotated using the STRINGdb package^53^ (using high-confidence interactions with a score > 400). Weighted joint probability (JP) heatmaps were constructed to visualize how protein–protein interactions (PPIs) are distributed across the abundance-rank space. For each protein-protein interaction annotated in the sorted matrix, we assigned a weight that combines the interaction confidence score with an additional boost for TP-TP pairs (and a smaller boost for TP-Other pairs), and then accumulated these weighted interactions into the corresponding abundance-rank bins. The total weighed counts in each bin were re-scaled to obtain probability-like values.

#### Volcano plot

Generated using dataframe3 and the ggplot2 package.

#### TP count percentage and abundance percentage plots

Compiled from the final protein lists and plotted using GraphPad Prism 9.

#### Precision-rank curve

Generated from the final protein lists, ranked by MS score in descending order. The TP set included all proteins with “SG/Nucleolus” in their annotation. Plotted using GraphPad Prism 9.

#### Euler diagram

Generated using the eulerr package and the final protein lists.

#### Protein sequence analysis

Local charge clustering ratio: (Number of adjacent amino acid pairs with the same charge) / (Total number of adjacent pairs with either the same or opposite charge).

Mean IDR length: Protein sequences were converted into a Boolean (T/F) vector indicating the presence of ten amino acids prone to disorder (A, R, G, Q, S, P, E, K, N, T). The mean length of continuous disordered segments was then calculated.

Fraction(charged): (|D| + |E| + |K| + |R| + |H|) / Total number of amino acids.

Poly-AA Freq: Frequency of four consecutive identical amino acids per 100 amino acids.

GSEA-like analysis of sequence features was performed using the clusterProfiler^87^ and GseaVis^88^ packages.Total protein list was generated by combining all detected 293T proteins from SG_batch 1 (4000-5000 proteins in intermediate protein list).

#### Phase-APEX2-seq data analysis

Phase-APEX2-Seq reads were trimmed for adapters and low-quality bases (-q 30 --paired) using Trim Galore (v0.6.7) and aligned to the human genome (T2T-CHM13v2) with Bowtie2. Alignments were sorted and indexed with samtools.

For visualization, BAM files were RPKM-normalized into BigWig files, and TSA-Seq enrichment scores were calculated as described^71,89^ using a 20 kb sliding window in Python. For the artificial rDNA reference (hs1-rDNA)^90^, a 2 kb sliding window was applied for higher-resolution inspection. Enrichment scores were quantified across different transcript biotypes based on annotations in GCF_009914755.1_T2T-CHM13v2.0_genomic.gtf and visualized as violin plots.

Peaks were called using MACS2 with merged input as control (--broad --broad-cutoff 0.1 -q 0.05 -f BAMPE). Downstream analyses were conducted in R, using the rtracklayer and GenomicRanges packages to evaluate peak counts and genomic distributions. Overlapping peaks were identified with the findOverlapsOfPeaks function from ChIPpeakAnno, and peak annotation was performed using annotatePeak. Gene Ontology (GO) enrichment analysis was carried out using enrichGO from the clusterProfiler package.

## REFERENCES

1. Brangwynne, C.P., Mitchison, T.J., and Hyman, A.A. (2011). Active liquid-like behavior of nucleoli determines their size and shape in *Xenopus laevis* oocytes. Proceedings of the National Academy of Sciences 108, 4334–4339.

2. Weber, S.C., and Brangwynne, C.P. (2015). Inverse Size Scaling of the Nucleolus by a Concentration-Dependent Phase Transition. Curr. Biol. 25, 641–646.

3. Feric, M., Vaidya, N., Harmon, T.S., Mitrea, D.M., Zhu, L., Richardson, T.M., Kriwacki, R.W., Pappu, R.V., and Brangwynne, C.P. (2016). Coexisting Liquid Phases Underlie Nucleolar Subcompartments. Cell 165, 1686–1697.

4. Wippich, F., Bodenmiller, B., Trajkovska, M.G., Wanka, S., Aebersold, R., and Pelkmans, L. (2013). Dual Specificity Kinase DYRK3 Couples Stress Granule Condensation/Dissolution to mTORC1 Signaling. Cell 152, 791–805.

5. Molliex, A., Temirov, J., Lee, J., Coughlin, M., Kanagaraj, A.P., Kim, H.J., Mittag, T., and Taylor, J.P. (2015). Phase Separation by Low Complexity Domains Promotes Stress Granule Assembly and Drives Pathological Fibrillization. Cell 163, 123–133.

6. Guillén-Boixet, J., Kopach, A., Holehouse, A.S., Wittmann, S., Jahnel, M., Schlüßler, R., Kim, K., Trussina, I.R.E.A., Wang, J., and Mateju, D., et al. (2020). RNA-Induced Conformational Switching and Clustering of G3BP Drive Stress Granule Assembly by Condensation. Cell 181, 346–361.

7. Yang, P., Mathieu, C., Kolaitis, R., Zhang, P., Messing, J., Yurtsever, U., Yang, Z., Wu, J., Li, Y., and Pan, Q., et al. (2020). G3BP1 Is a Tunable Switch that Triggers Phase Separation to Assemble Stress Granules. Cell 181, 325–345.

8. Mackenzie, I.R., Nicholson, A.M., Sarkar, M., Messing, J., Purice, M.D., Pottier, C., Annu, K., Baker, M., Perkerson, R.B., and Kurti, A., et al. (2017). TIA1 Mutations in Amyotrophic Lateral Sclerosis and Frontotemporal Dementia Promote Phase Separation and Alter Stress Granule Dynamics. Neuron 95, 808–816.

9. Gao, Y., Li, X., Li, P., and Lin, Y. (2022). A brief guideline for studies of phase-separated biomolecular condensates. Nat. Chem. Biol. 18, 1307–1318.

10. Hung, V., Udeshi, N.D., Lam, S.S., Loh, K.H., Cox, K.J., Pedram, K., Carr, S.A., and Ting, A.Y. (2016). Spatially resolved proteomic mapping in living cells with the engineered peroxidase APEX2. Nat. Protoc. 11, 456–475.

11. Qin, W., Cho, K.F., Cavanagh, P.E., and Ting, A.Y. (2021). Deciphering molecular interactions by proximity labeling. Nat. Methods 18, 133–143.

12. Zhang, S., Tang, Q., Zhang, X., and Chen, X. (2024). Proximitomics by Reactive Species. ACS Cent. Sci. 10, 1135–1147.

13. Irgen-Gioro, S., Yoshida, S., Walling, V., and Chong, S. (2022). Fixation can change the appearance of phase separation in living cells. Elife 11, e79903.

14. Lyon, A.S., Peeples, W.B., and Rosen, M.K. (2021). A framework for understanding the functions of biomolecular condensates across scales. Nat. Rev. Mol. Cell Biol. 22, 215–235.

15. Alberti, S., Gladfelter, A., and Mittag, T. (2019). Considerations and Challenges in Studying Liquid-Liquid Phase Separation and Biomolecular Condensates. Cell 176, 419–434.

16. Thul, P.J., åkesson, L., Wiking, M., Mahdessian, D., Geladaki, A., Ait Blal, H., Alm, T., Asplund, A., Björk, L., and Breckels, L.M., et al. (2017). A subcellular map of the human proteome. Science 356, eaal3321.

17. Brignole, C., Bensa, V., Fonseca, N.A., Del Zotto, G., Bruno, S., Cruz, A.F., Malaguti, F., Carlini, B., Morandi, F., and Calarco, E., et al. (2021). Cell surface Nucleolin represents a novel cellular target for neuroblastoma therapy. J. Exp. Clin. Cancer Res. 40, 180.

18. Perr, J., Langen, A., Almahayni, K., Nestola, G., Chai, P., Lebedenko, C.G., Volk, R.F., Detrés, D., Caldwell, R.M., and Spiekermann, M., et al. (2025). RNA-binding proteins and glycoRNAs form domains on the cell surface for cell-penetrating peptide entry. Cell 188, 1878–1895.

19. Hovanessian, A.G., Puvion-Dutilleul, F., Nisole, S., Svab, J., Perret, E., Deng, J.S., and Krust, B. (2000). The cell-surface-expressed nucleolin is associated with the actin cytoskeleton. Exp. Cell Res. 261, 312–328.

20. Qin, W., Cheah, J.S., Xu, C., Messing, J., Freibaum, B.D., Boeynaems, S., Taylor, J.P., Udeshi, N.D., Carr, S.A., and Ting, A.Y. (2023). Dynamic mapping of proteome trafficking within and between living cells by TransitID. Cell 186, 3307–3324.

21. Xing, W., Muhlrad, D., Parker, R., and Rosen, M.K. (2020). A quantitative inventory of yeast P body proteins reveals principles of composition and specificity. Elife 9, e56525.

22. Keber, F.C., Nguyen, T., Mariossi, A., Brangwynne, C.P., and Wühr, M. (2024). Evidence for widespread cytoplasmic structuring into mesoscale condensates. Nat. Cell Biol. 26, 346–352.

23. Lam, S.S., Martell, J.D., Kamer, K.J., Deerinck, T.J., Ellisman, M.H., Mootha, V.K., and Ting, A.Y. (2015). Directed evolution of APEX2 for electron microscopy and proximity labeling. Nat. Methods 12, 51–54.

24. Branon, T.C., Bosch, J.A., Sanchez, A.D., Udeshi, N.D., Svinkina, T., Carr, S.A., Feldman, J.L., Perrimon, N., and Ting, A.Y. (2018). Efficient proximity labeling in living cells and organisms with TurboID. Nat. Biotechnol. 36, 880–887.

25. Seath, C.P., Burton, A.J., Sun, X., Lee, G., Kleiner, R.E., Macmillan, D.W.C., and Muir, T.W. (2023). Tracking chromatin state changes using nanoscale photo-proximity labelling. Nature 616, 574–580.

26. Pan, C., Knutson, S.D., Huth, S.W., and Macmillan, D.W.C. (2025). µMap proximity labeling in living cells reveals stress granule disassembly mechanisms. Nat. Chem. Biol. 21, 490–500.

27. Lin, Z., Schaefer, K., Lui, I., Yao, Z., Fossati, A., Swaney, D.L., Palar, A., Sali, A., and Wells, J.A. (2024). Multiscale photocatalytic proximity labeling reveals cell surface neighbors on and between cells. Science 385, eadl5763.

28. Lee, C., Quintana, A., Suppanz, I., Gomez-Auli, A., Mittler, G., and Cissé, I.I. (2024). Light-induced targeting enables proteomics on endogenous condensates. Cell 187, 7079–7090.

29. Li, K., Xie, X., Gao, R., Chen, Z., Yang, M., Wen, Z., Weng, Y., Fan, X., Zhang, G., and Liu, L., et al. (2024). Spatiotemporal protein interactome profiling through condensation-enhanced photocrosslinking. Nat. Chem.

30. Lee, S., Cheah, J.S., Zhao, B., Xu, C., Roh, H., Kim, C.K., Cho, K.F., Udeshi, N.D., Carr, S.A., and Ting, A.Y. (2023). Engineered allostery in light-regulated LOV-Turbo enables precise spatiotemporal control of proximity labeling in living cells. Nat. Methods 20, 908–917.

31. Han, Y., Branon, T.C., Martell, J.D., Boassa, D., Shechner, D., Ellisman, M.H., and Ting, A. (2019). Directed Evolution of Split APEX2 Peroxidase. ACS Chem. Biol. 14, 619–635.

32. Cho, K.F., Branon, T.C., Rajeev, S., Svinkina, T., Udeshi, N.D., Thoudam, T., Kwak, C., Rhee, H.W., Lee, I.K., and Carr, S.A., et al. (2020). Split-TurboID enables contact-dependent proximity labeling in cells. Proc. Natl. Acad. Sci. U. S. A. 117, 12143–12154.

33. Qu, D., Li, Y., Liu, Q., Cao, B., Cao, M., Lin, X., Shen, C., Zou, P., Zhou, H., and Zhang, W., et al. (2025). Photoactivated SOPP3 enables APEX2-mediated proximity labeling with high spatio-temporal resolution in live cells. Cell Res. 35, 149–152.

34. Sroka, T.J., Sanwald, L.K., Prasai, A., Hoeren, J., von der Malsburg, K., Chaumet, V., Haberkant, P., Feistel, K., and Mick, D.U. (2025). iAPEX: Improved APEX-based proximity labeling for subcellular proteomics using an enzymatic reaction cascade. bioRxiv, 2021-2025.

35. Shan, L., Xu, G., Yao, R., Luan, P., Huang, Y., Zhang, P., Pan, Y., Zhang, L., Gao, X., and Li, Y., et al. (2023). Nucleolar URB1 ensures 3′ ETS rRNA removal to prevent exosome surveillance. Nature 615, 526–534.

36. Go, C.D., Knight, J.D.R., Rajasekharan, A., Rathod, B., Hesketh, G.G., Abe, K.T., Youn, J., Samavarchi-Tehrani, P., Zhang, H., and Zhu, L.Y., et al. (2021). A proximity-dependent biotinylation map of a human cell. Nature 595, 120–124.

37. Yap, K., Chung, T.H., and Makeyev, E.V. (2022). Hybridization-proximity labeling reveals spatially ordered interactions of nuclear RNA compartments. Mol. Cell 82, 463–478.

38. Kawai, K., Hirayama, T., Imai, H., Murakami, T., Inden, M., Hozumi, I., and Nagasawa, H. (2022). Molecular Imaging of Labile Heme in Living Cells Using a Small Molecule Fluorescent Probe. J. Am. Chem. Soc. 144, 3793–3803.

39. Hanna, D.A., Harvey, R.M., Martinez-Guzman, O., Yuan, X., Chandrasekharan, B., Raju, G., Outten, F.W., Hamza, I., and Reddi, A.R. (2016). Heme dynamics and trafficking factors revealed by genetically encoded fluorescent heme sensors. Proceedings of the National Academy of Sciences 113, 7539–7544.

40. Atamna, H., Brahmbhatt, M., Atamna, W., Shanower, G.A., and Dhahbi, J.M. (2015). ApoHRP-based assay to measure intracellular regulatory heme. Metallomics 7, 309–321.

41. Torra, J., Lafaye, C., Signor, L., Aumonier, S., Flors, C., Shu, X., Nonell, S., Gotthard, G., and Royant, A. (2019). Tailing miniSOG: structural bases of the complex photophysics of a flavin-binding singlet oxygen photosensitizing protein. Sci. Rep. 9.

42. Barnett, M.E., Baran, T.M., Foster, T.H., and Wojtovich, A.P. (2018). Quantification of light-induced miniSOG superoxide production using the selective marker, 2-hydroxyethidium. Free. Radic. Biol. Med. 116, 134–140.

43. Somwar, R., Erdjument-Bromage, H., Larsson, E., Shum, D., Lockwood, W.W., Yang, G., Sander, C., Ouerfelli, O., Tempst, P.J., and Djaballah, H., et al. (2011). Superoxide dismutase 1 (SOD1) is a target for a small molecule identified in a screen for inhibitors of the growth of lung adenocarcinoma cell lines. Proceedings of the National Academy of Sciences 108, 16375–16380.

44. Shu, X., Lev-Ram, V., Deerinck, T.J., Qi, Y., Ramko, E.B., Davidson, M.W., Jin, Y., Ellisman, M.H., and Tsien, R.Y. (2011). A Genetically Encoded Tag for Correlated Light and Electron Microscopy of Intact Cells, Tissues, and Organisms. PLoS Biol. 9, e1001041.

45. Lafaye, C., Aumonier, S., Torra, J., Signor, L., von Stetten, D., Noirclerc-Savoye, M., Shu, X., Ruiz-González, R., Gotthard, G., and Royant, A., et al. (2022). Riboflavin-binding proteins for singlet oxygen production. Photochem. Photobiol. Sci. 21, 1545–1555.

46. Westberg, M., Bregnhøj, M., Etzerodt, M., and Ogilby, P.R. (2017). No Photon Wasted: An Efficient and Selective Singlet Oxygen Photosensitizing Protein. The Journal of Physical Chemistry B 121, 9366–9371.

47. Zhai, Y., Huang, X., Zhang, K., Huang, Y., Jiang, Y., Cui, J., Zhang, Z., Chiu, C.K.C., Zhong, W., and Li, G. (2022). Spatiotemporal-resolved protein networks profiling with photoactivation dependent proximity labeling. Nat. Commun. 13, 4906.

48. Ren, Z., Tang, W., Peng, L., and Zou, P. (2023). Profiling stress-triggered RNA condensation with photocatalytic proximity labeling. Nat. Commun. 14, 7390.

49. Hananya, N., Ye, X., Koren, S., and Muir, T.W. (2023). A genetically encoded photoproximity labeling approach for mapping protein territories. Proceedings of the National Academy of Sciences 120, e2075628176.

50. Zhou, Y., Wang, G., Wang, P., Li, Z., Yue, T., Wang, J., and Zou, P. (2019). Expanding APEX2 Substrates for Proximity-Dependent Labeling of Nucleic Acids and Proteins in Living Cells. Angewandte Chemie 131, 11889–11893.

51. Marmor-Kollet, H., Siany, A., Kedersha, N., Knafo, N., Rivkin, N., Danino, Y.M., Moens, T.G., Olender, T., Sheban, D., and Cohen, N., et al. (2020). Spatiotemporal Proteomic Analysis of Stress Granule Disassembly Using APEX Reveals Regulation by SUMOylation and Links to ALS Pathogenesis. Mol. Cell 80, 876–891.

52. Li, P., Banjade, S., Cheng, H., Kim, S., Chen, B., Guo, L., Llaguno, M., Hollingsworth, J.V., King, D.S., and Banani, S.F., et al. (2012). Phase transitions in the assembly of multivalent signalling proteins. Nature 483, 336–340.

53. Szklarczyk, D., Kirsch, R., Koutrouli, M., Nastou, K., Mehryary, F., Hachilif, R., Gable, A.L., Fang, T., Doncheva, N.T., and Pyysalo, S., et al. (2023). The STRING database in 2023: protein–protein association networks and functional enrichment analyses for any sequenced genome of interest. Nucleic. Acids. Res. 51, D638–D646.

54. Pei, G., Wang, X., Geng, D., Chen, Z., Xu, W., Li, T., and Li, P. (2024). Sequence Composition Dictates Condensate Miscibility. bioRxiv, 2011-2024.

55. Ruff, K.M., King, M.R., Ying, A.W., Liu, V., Pant, A., Lieberman, W.E., Shinn, M.K., Su, X., Kadoch, C., and Pappu, R.V. (2025). Molecular grammars of intrinsically disordered regions that span the human proteome. bioRxiv, 2022-2025.

56. Lyons, H., Veettil, R.T., Pradhan, P., Fornero, C., De La Cruz, N., Ito, K., Eppert, M., Roeder, R.G., and Sabari, B.R. (2023). Functional partitioning of transcriptional regulators by patterned charge blocks. Cell 186, 327–345.

57. Goldstrohm, A.C., Hall, T.M.T., and Mckenney, K.M. (2018). Post-transcriptional Regulatory Functions of Mammalian Pumilio Proteins. Trends Genet. 34, 972–990.

58. Vessey, J.P., Vaccani, A., Xie, Y., Dahm, R., Karra, D., Kiebler, M.A., and Macchi, P. (2006). Dendritic Localization of the Translational Repressor Pumilio 2 and Its Contribution to Dendritic Stress Granules. J. Neurosci. 26, 6496–6508.

59. Majerciak, V., Zhou, T., Kruhlak, M.J., and Zheng, Z. (2023). RNA helicase DDX6 and scaffold protein GW182 in P-bodies promote biogenesis of stress granules. Nucleic. Acids. Res. 51, 9337–9355.

60. Kaehler, C., Isensee, J., Nonhoff, U., Terrey, M., Hucho, T., Lehrach, H., and Krobitsch, S. (2012). Ataxin-2-Like Is a Regulator of Stress Granules and Processing Bodies. PLoS One 7, e50134.

61. Elden, A.C., Kim, H., Hart, M.P., Chen-Plotkin, A.S., Johnson, B.S., Fang, X., Armakola, M., Geser, F., Greene, R., and Lu, M.M., et al. (2010). Ataxin-2 intermediate-length polyglutamine expansions are associated with increased risk for ALS. Nature 466, 1069–1075.

62. Fu, Y., and Zhuang, X. (2020). m6A-binding YTHDF proteins promote stress granule formation. Nat. Chem. Biol. 16, 955–963.

63. Krüger, D.M., Neubacher, S., and Grossmann, T.N. (2018). Protein–RNA interactions: structural characteristics and hotspot amino acids. RNA 24, 1457–1465.

64. Requião, R.D., Fernandes, L., de Souza, H.J.A., Rossetto, S., Domitrovic, T., and Palhano, F.L. (2017). Protein charge distribution in proteomes and its impact on translation. PLoS Comput. Biol. 13, e1005549.

65. Paloni, M., Bussi, G., and Barducci, A. (2021). Arginine multivalency stabilizes protein/RNA condensates. Protein. Sci. 30, 1418–1426.

66. Wang, J., Choi, J., Holehouse, A.S., Lee, H.O., Zhang, X., Jahnel, M., Maharana, S., Lemaitre, R., Pozniakovsky, A., and Drechsel, D., et al. (2018). A Molecular Grammar Governing the Driving Forces for Phase Separation of Prion-like RNA Binding Proteins. Cell 174, 688–699.

67. Kar, M., Posey, A.E., Dar, F., Hyman, A.A., and Pappu, R.V. (2021). Glycine-Rich Peptides from FUS Have an Intrinsic Ability to Self-Assemble into Fibers and Networked Fibrils. Biochemistry. 60, 3213–3222.

68. Yuan, L., Mao, L., Huang, Y., Outeiro, T.F., Li, W., Vieira, T.C.R.G., and Li, J. (2025). Stress granules: emerging players in neurodegenerative diseases. Translational Neurodegeneration 14, 22.

69. Zhang, X., Vigers, M., Mccarty, J., Rauch, J.N., Fredrickson, G.H., Wilson, M.Z., Shea, J., Han, S., and Kosik, K.S. (2020). The proline-rich domain promotes Tau liquid–liquid phase separation in cells. J. Cell. Biol. 219, e202006054.

70. Lieberman, A.P., Shakkottai, V.G., and Albin, R.L. (2019). Polyglutamine Repeats in Neurodegenerative Diseases. Annual review of pathology 14, 1–27.

71. Chen, Y., Zhang, Y., Wang, Y., Zhang, L., Brinkman, E.K., Adam, S.A., Goldman, R., van Steensel, B., Ma, J., and Belmont, A.S. (2018). Mapping 3D genome organization relative to nuclear compartments using TSA-Seq as a cytological ruler. J. Cell. Biol. 217, 4025–4048.

72. Ke, M., Yuan, X., He, A., Yu, P., Chen, W., Shi, Y., Hunter, T., Zou, P., and Tian, R. (2021). Spatiotemporal profiling of cytosolic signaling complexes in living cells by selective proximity proteomics. Nat. Commun. 12, 71.

73. Hori, Y., Shimamoto, A., and Kobayashi, T. (2021). The human ribosomal DNA array is composed of highly homogenized tandem clusters. Genome Res. 31, 1971–1982.

74. Gucwa, M., and Ikegami, K. (2025). Cell-cycle dynamics of nascent transcription and mature RNA accumulation are concordant in normal fibroblasts. bioRxiv, 2025-2029.

75. Liu, Y., Chen, S., Wang, S., Soares, F., Fischer, M., Meng, F., Du, Z., Lin, C., Meyer, C., and Decaprio, J.A., et al. (2017). Transcriptional landscape of the human cell cycle. Proceedings of the National Academy of Sciences 114, 3473–3478.

76. Palozola, K.C., Donahue, G., Liu, H., Grant, G.R., Becker, J.S., Cote, A., Yu, H., Raj, A., and Zaret, K.S. (2017). Mitotic transcription and waves of gene reactivation during mitotic exit. Science 358, 119–122.

77. Bersaglieri, C., Kresoja-Rakic, J., Gupta, S., Bär, D., Kuzyakiv, R., Panatta, M., and Santoro, R. (2022). Genome-wide maps of nucleolus interactions reveal distinct layers of repressive chromatin domains. Nat. Commun. 13, 1483.

78. Walavalkar, K., Gupta, S., Kresoja-Rakic, J., Raingeval, M., Mungo, C., and Santoro, R. (2025). Single-cell dynamics of genome-nucleolus interactions captured by nucleolar laser microdissection (NoLMseq). bioRxiv, 2024-2026.

79. Fazal, F.M., Han, S., Parker, K.R., Kaewsapsak, P., Xu, J., Boettiger, A.N., Chang, H.Y., and Ting, A.Y. (2019). Atlas of Subcellular RNA Localization Revealed by APEX-Seq. Cell 178, 473–490.

80. Sun, X., Zhang, Y., Lu, W., Guo, H., He, G., Luo, S., Guo, H., Zhang, Z., Wang, W., and Chu, L., et al. (2025). Precise and*In Vivo* -Compatible Spatial Proteomics via Bioluminescence-Triggered Photocatalytic Proximity Labeling. ACS Cent. Sci. 11, 1611–1626.

81. Fragpipe tutorial. https://fragpipe.nesvilab.org/docs/tutorial_fragpipe.html.

82. Yu, F., Haynes, S.E., and Nesvizhskii, A.I. (2021). IonQuant Enables Accurate and Sensitive Label-Free Quantification With FDR-Controlled Match-Between-Runs. Molecular & Cellular Proteomics 20.

83. Rath, S., Sharma, R., Gupta, R., Ast, T., Chan, C., Durham, T.J., Goodman, R.P., Grabarek, Z., Haas, M.E., and Hung, W.H.W., et al. (2021). MitoCarta3.0: an updated mitochondrial proteome now with sub-organelle localization and pathway annotations. Nucleic. Acids. Res. 49, D1541–D1547.

84. Carbon, S., Ireland, A., Mungall, C.J., Shu, S., Marshall, B., Lewis, S., The, A.H., and The, W.P.W.G. (2009). AmiGO: online access to ontology and annotation data. Bioinformatics 25, 288–289.

85. Robin, X., Turck, N., Hainard, A., Tiberti, N., Lisacek, F., Sanchez, J., and Müller, M. (2011). pROC: an open-source package for R and S+ to analyze and compare ROC curves. BMC Bioinformatics 12, 77.

86. Ritchie, M.E., Phipson, B., Wu, D., Hu, Y., Law, C.W., Shi, W., and Smyth, G.K. (2015). limma powers differential expression analyses for RNA-sequencing and microarray studies. Nucleic. Acids. Res. 43, e47.

87. Xu, S., Hu, E., Cai, Y., Xie, Z., Luo, X., Zhan, L., Tang, W., Wang, Q., Liu, B., and Wang, R., et al. (2024). Using clusterProfiler to characterize multiomics data. Nat. Protoc. 19, 3292–3320.

88. Zhang, J., Li, H., Tao, W., and Zhou, J. (2025). GseaVis: An R Package for Enhanced Visualization of Gene Set Enrichment Analysis in Biomedicine. Med Research 1, 131–135.

89. Zhang, L., Zhang, Y., Chen, Y., Gholamalamdari, O., Wang, Y., Ma, J., and Belmont, A.S. (2021). TSA-seq reveals a largely conserved genome organization relative to nuclear speckles with small position changes tightly correlated with gene expression changes. Genome Res. 31, 251–264.

90. George, S.S., Pimkin, M., and Paralkar, V.R. (2023). Construction and validation of customized genomes for human and mouse ribosomal DNA mapping. Journal of Biological Chemistry 299, 104766.

